# Multi-view deep learning of highly multiplexed imaging data improves association of cell states with clinical outcomes

**DOI:** 10.1101/2025.03.14.643377

**Authors:** Shanza Ayub, Hartland W. Jackson, Alina Selega, Kieran R. Campbell

## Abstract

Highly multiplexed imaging technologies can quantify the expression of 10s-100s of proteins and their modifications at sub-cellular resolution. Applications to tissue specimens across a broad array of pathologies have enabled patient subtyping in terms of novel cell states and spatial cell communities and linked their prevalence and overall architecture to patient outcomes. While highly multiplexed imaging analysis workflows typically summarize each cell in terms of its post-segmentation mean expression, additional cellular information can be quantified including cell morphology, sub-cellular expression patterns, and spatial cellular context, ultimately giving a multi-modal view of each cell. While deep latent variable models such as variational autoencoders (VAEs) are well established for other multi-modal single-cell assays, their ability to integrate these multiple views of a cell from highly multiplexed imaging data remains largely unknown.

Here, we explore the abilities of multi-modal VAEs to learn unified latent cellular representations from multiple views of each single-cell quantified from highly multiplexed imaging, including mean expression, morphology, sub-cellular protein co-localization, and spatial cellular context, while conditioning on technical and batch specific effects. We first discuss strategies for training and hyperparameter optimization of such models across a set of highly multiplexed imaging datasets of breast and melanoma cancers. We next quantify the relevance of the integrated multi-modal latent space in predicting patient-specific clinical outcomes and demonstrate competitive performance across a range of outcomes compared to existing baselines. Then, we explore the ability of the cellular representations to learn cellular phenotypes that align with known cell types and find that the expression-specific latent representation can identify clusters more closely aligned with known lymphocyte and epithelial subpopulations compared to established workflows. Finally, we test the ability of each representation to perform consistent batch integration across different cohorts and discuss the considerations on taking known background variables into account.

## Introduction

Highly multiplexed imaging (HMI) technologies can quantify the expression of 10-100 proteins and their modifications within single cells while retaining their spatial locations. These include immunofluorescence based assays such as cyclic immunofluorescence which use optical imaging^1,2^, and mass cytometry based assays such as Imaging Mass Cytometry (IMC) and Multiplexed Ion Beam Imaging^3,4^. These technologies have revolutionized cancer research by elucidating spatial tumour heterogeneity with significant consequences for patient outcomes^5,6^. The standard analytical workflow of HMI data includes cell segmentation that identifies single cells within the image and generates a cell segmentation mask, containing cell IDs and their spatial locations. Most studies subsequently cluster cells based on the mean protein expression and use spatial information contained in these images to detect patterns in tissue architecture^2^. However, cellular states and functions are additionally determined by protein subcellular localization, cellular morphology, and the influence of local spatial architecture within a tissue^7–9^.

Deep learning methods are gaining popularity as analysis tools for HMI data but have been largely limited to image segmentation and cell type annotation^10^. Improved segmentation using deep learning in HMI data allows for accurate identification of cells within a tissue image which is crucial for cell type assignment and further downstream analysis has revealed distinct communities in the tumour microenvironment (TME) associated with clinical outcomes^11–13^. Deep learning methods have also improved and automated identification of major cell types in tissues^14–16^, but these methods cannot be used to identify variations within these major cell types as they are limited to using single-cell protein expression alone and do not take into account the spatial context of the cells. Some studies incorporate spatial information and the tissue context to understand cell states^17,18^ but these ignore the impacts of protein subcellular localization on cell state and differences in cell morphology.

Spatial transcriptomics (ST) and single-cell RNA sequencing (scRNA-seq) have seen more application of deep learning methods, largely for batch correction and spot deconvolution with the aim to identify cell types and states^19–21^. Further, existing deep learning methods integrate different modalities such as totalVI, which integrates cell surface protein expression with transcriptomics data^22^ and DeepST and SpatialDIVA which use data from histology and spatial transcriptomics images to identify spatial domains of clinical importance^23,24^. Many of these models follow a Variational Autoencoder (VAE) framework and have shown state-of-the-art performance in integrating multi-modal data inputs, enabling the classification of cell states not identifiable by a single data type alone^25^. VAEs are generative deep unsupervised models that learn a latent space—a compressed lower dimensional representation of the high dimensional input data—that tries to strike a balance between accurate data reconstruction and model generalizability^26–28^. The lower dimensional latent representation learnt by a VAE captures relevant features in the data that can be used for downstream analysis such as clustering and cell state identification as well as associations with clinical variables using patient data^29,30^.

Here, we propose a multi-modal VAE for highly multiplexed imaging data (hmiVAE) that learns combined and mode-specific latent representations from input mean expression, morphology, protein nuclear co-localization, and spatial cellular context, while conditioning on technical and batch specific effects. We apply this model to several IMC datasets of breast and melanoma cancers^5,6,31^ and quantify the extent to which the integrated multi-modal latent space is predictive of specific clinical outcomes compared to existing approaches. Next, we show that our expression-specific representation can identify cell clusters more closely aligned with known lymphocyte and epithelial subpopulations compared to established workflows. Finally, we test the ability of our model to perform consistent batch integration across different cohorts. Together, this study provides the first exploration of using multi-modal VAEs to learn meaningful cell representations from highly multiplexed imaging data and demonstrates its utility on several common downstream analysis tasks.

## Results

### Exploring hyperparameter effects and unveiling cellular heterogeneity using multi-view integration with hmiVAE

We first developed a data processing pipeline to extract four views from IMC data: (i) mean protein expression, (ii) protein nuclear co-localization, (iii) cell morphology and (iv) cell spatial context. We extract these views for each cell within an image. First, we extract mean protein cellular expression by taking the mean of the protein expression across all the pixels belonging to a cell. Second, we use the per-pixel protein expression counts and find the correlation between these and the mean nuclear score as a proxy for nuclear co-localization. Third, we compute the cell morphology features using image processing libraries in Python (**Methods**). Finally, we summarise the expression counts, nuclear co-localization, and morphology for the *k* nearest neighbours of each cell to get the spatial context. Furthermore, highly multiplexed imaging data suffers from technical artefacts such as speckles resulting from non-specific antibody binding and uneven signal intensities across samples^32^. To control for such batch effects, we condition on sample specific identifiers and likely technical artefacts by including a one-hot encoding of cells belonging to each sample and computing covariates such as per-sample mean image intensity (**Methods**). These views and covariates form the input to our highly multiplexed imaging variational autoencoder (hmiVAE). hmiVAE builds on a standard VAE architecture consisting of two neural networks, an encoder and decoder. The encoder takes in each view (expression, nuclear co-localization, morphology and spatial context) and batch-specific information to first learn a separate representation for each view and then concatenates them to learn an integrated latent space. The decoder then samples from this integrated latent space to reconstruct each view (**Fig. 1A & Methods**). The integrated latent space, as well as the view-specific embeddings can then be used for downstream tasks such as cell type interpretation and associations with clinical variables.

**Figure 1.**
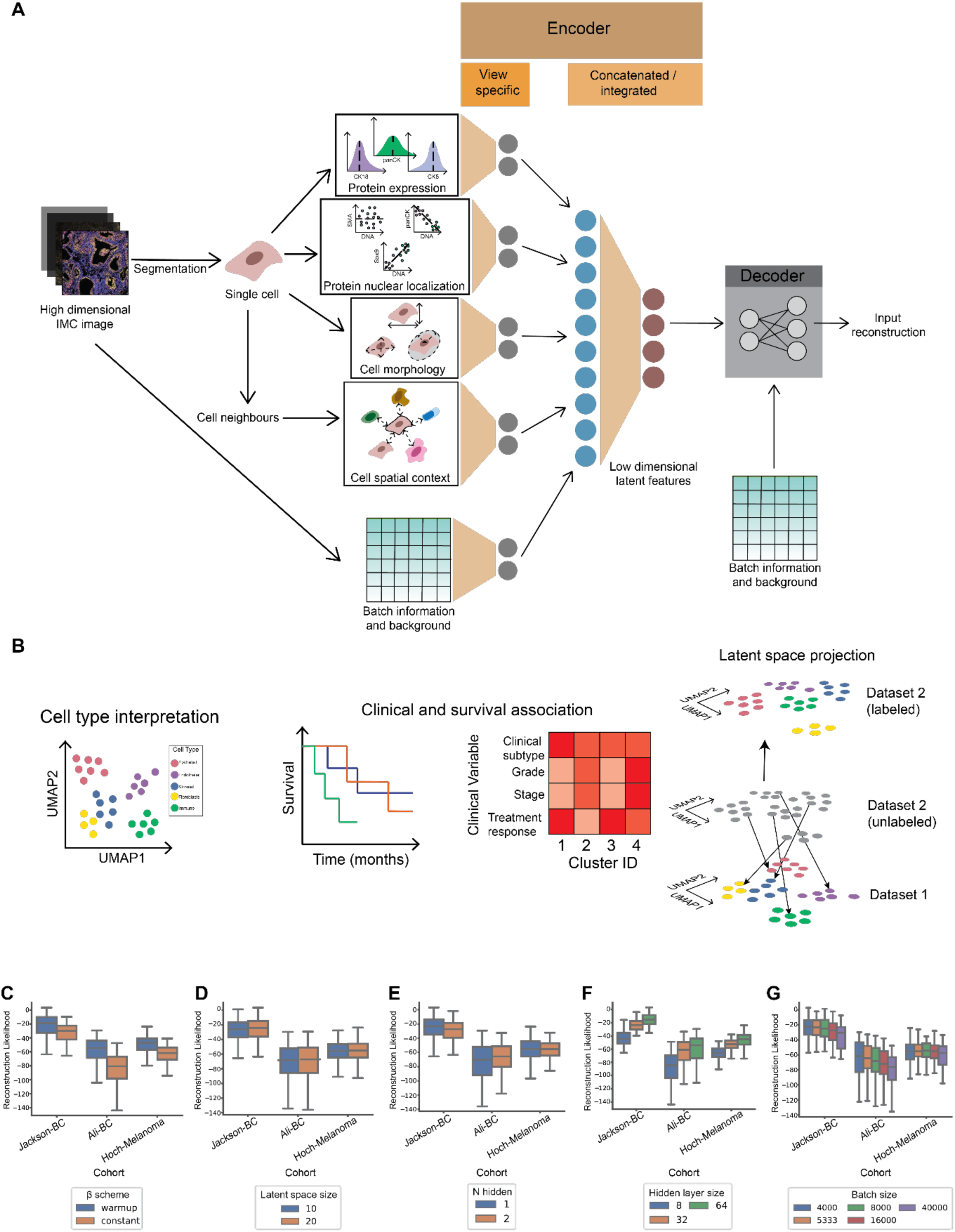
Multi-view integration of highly multiplexed imaging data using multi-modal variational autoencoders. **A** hmiVAE architecture. hmiVAE takes in inputs from 4 cell-specific views: protein expression, protein-nuclear co-localization, cell morphology and cell spatial context. At the encoder stage, it first learns separate embeddings for each view which are then concatenated to learn a combined latent space. The decoder samples from this latent space to reconstruct the inputs. **B** Downstream tasks. The latent space and view-specific embeddings can be used for tasks such as cell type interpretation and association with clinical variables. We can also project cells from different datasets into the learnt latent space to find representations for these cells. **C - G** Hyperparameter tuning. Hyperparameter tuning results across datasets for beta scheme for KL-Divergence weighting (constant or warm-up) **(C),** latent space size **(D)**, number of hidden layers **(E),** size of hidden layers **(F)**, and batch size **(G)**.

As tuning hyperparameters is a key aspect of successfully training any deep learning method^33^,we considered how varying a set of parameters affected the validation loss. Specifically, we selected (i) the number of hidden layers, (ii) the size of the hidden layers (i.e. hidden layer dimension), (iii) latent space dimensionality, (iv) batch size, and (v) the weight of the KL divergence term with a warmup and constant schedule (**Methods**) and applied grid search over these to test how each impacted the reconstruction likelihood. We found that implementing a schedule on the weight resulted in better reconstruction than keeping it constant throughout training (**Fig. 1C**), a result also observed in other studies^22,34^. We also noticed that while the dimensionality of the latent space and the number of hidden layers did not have a large effect on the reconstruction likelihood (**Fig. 1D&E**), a larger hidden layer size resulted in better reconstruction (**Fig. 1F**). Similarly, having a smaller batch size during training also resulted in better reconstruction (**Fig. 1G**). We trained the models with the best-performing hyperparameters for each dataset, *Jackson-BC* and *Ali-BC* which are breast cancer IMC datasets from Jackson et al. (2020)^5^ and Ali et al. (2020)^6^ respectively with epithelial and stromal marker panel, and Hoch-Melanoma which is a melanoma IMC dataset from Hoch et al.(2022)^31^ with an immune marker panel (**Methods, Supplementary Table S1**). The resulting latent spaces from hmiVAE can be used for downstream analysis^19,22,23^. Using leiden clustering from the single-cell analysis Python library, scanpy^35^ (**Methods**), we clustered the resulting latent spaces, we refer to these as *integrated clusters* below.

To quantify the influence of each view on the resulting cell clusters, we individually ranked the features corresponding to each view (**Methods**) by the magnitude of their association with each integrated cluster (**Methods**). Visualizing the top three features per cluster shows that for each dataset many clusters had similar average expression profiles but differed in the localization of different proteins or the context of their neighbouring cells (**Fig. 2**). In the Ali-BC dataset (**Fig. 2B**), integrated clusters 12 and 21 both represent stromal cell clusters based on expression of Vimentin and Smooth muscle actin (SMA). However, they differ in terms of their spatial context: cluster 12 indicates stromal cells surrounded by apoptotic epithelial cells, given by cluster 12’s association with the expression of cPARP/cCasp3 and EpCAM in the neighbourhood, whereas cells in cluster 21 show a negative association with neighbourhood expression of cPARP/cCasp3 and EpCAM. Similarly, in the Hoch-Melanoma dataset, integrated clusters 4, 9 and 15 represent tumour cells, with cells in clusters in 9 and 15 having atypical morphology. Cluster 4 differs from others by the presence of a tumour suppressive environment indicated by the neighbourhood expression of markers for T regulatory cells (CD3, FOXP3 and TOX1) and macrophages (CD11b) (**Fig. 2C**). In the Jackson-BC dataset, we find integrated clusters 1, 7 and 5, which represent epithelial cells as indicated by expression but cells in cluster 1 differ in the self and neighbourhood expression of the hypoxia marker CAIX and Epidermal growth factor receptor (EGFR) as well as the nuclear co-localization of these markers (**Fig. 2A**).

**Figure 2.**
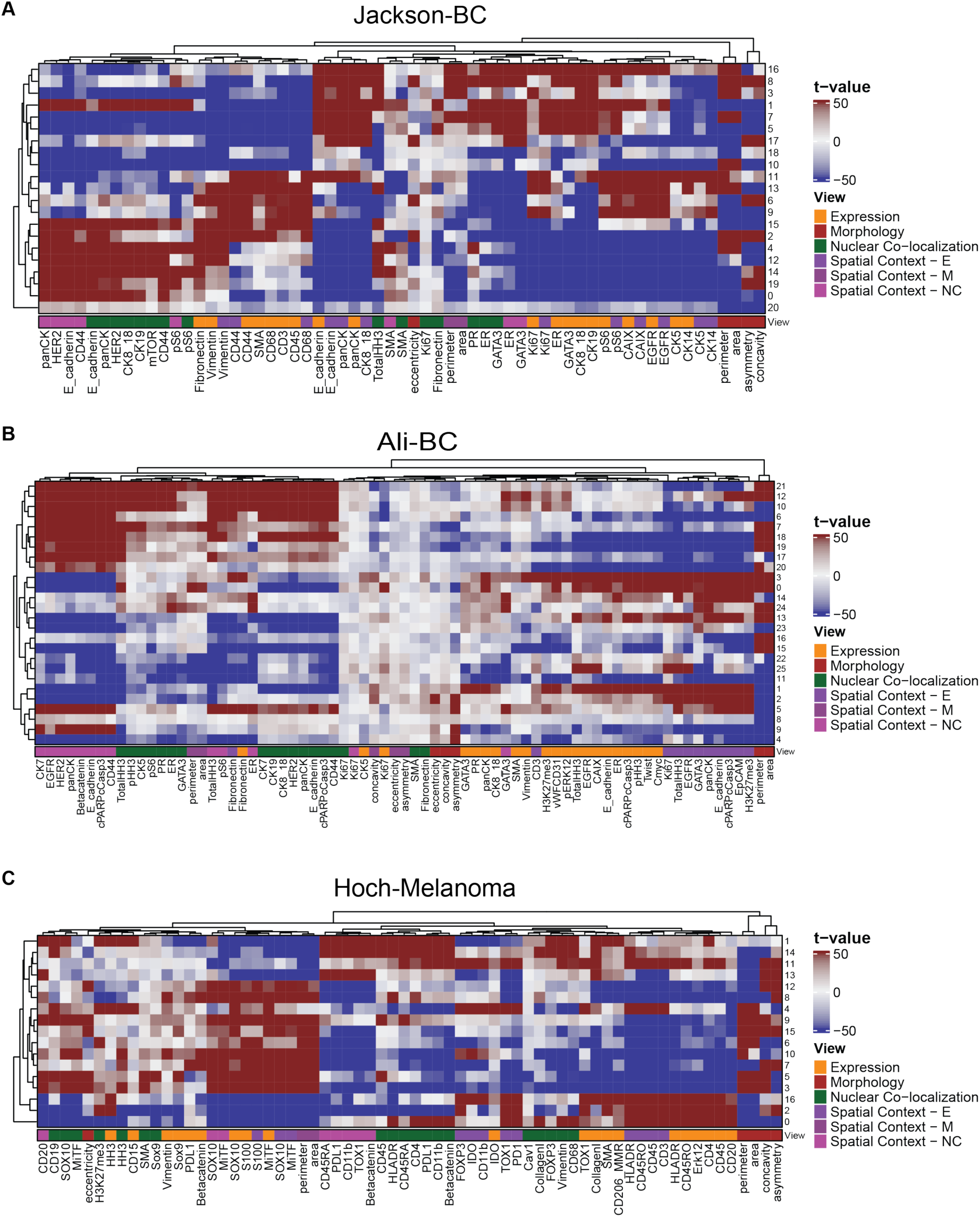
Ranking features across each view for integrated space clusters. Heatmaps showing the t-values for each feature representing their association with each integrated cluster in the dataset. Clusters in each dataset show differing patterns of within-cell protein expression, nuclear colocalization and morphology, and neighbourhood-cell protein expression, nuclear colocalization and morphology. **A** Association of clusters found from the integrated latent space of hmiVAE run on Jackson-BC. **B** Association of clusters found from the integrated latent space of hmiVAE run on Ali-BC. **C** Association of clusters found from the integrated latent space of hmiVAE run on Hoch-Melanoma. Spatial context - E (neighbourhood-cell protein expression), Spatial context - NC (neighbourhood-cell protein nuclear co-localization), Spatial context - M (neighbourhood-cell morphology).

### Expression-only embeddings improve cell type interpretation

Cell type annotation represents an important step in IMC data analysis workflow to provide insights into the cellular compositions of the tissues. Since this is commonly performed using the mean expression of proteins, we aimed to evaluate the efficacy of using the expression-specific cell representations from hmiVAE for this task and compared it to cell types found by clustering over the mean protein expression values using a standard community detection algorithm, Louvain^36^ and a popular clustering method, FlowSOM^37^ (**Methods**).

For the Ali-BC and Jackson-BC datasets that contain markers for major cell type lineages within epithelial, stromal and immune compartments, hmiVAE identified distinct clusters corresponding to CD3+ T cells, CD20+ B cells and CD68+ macrophages (**Fig. 3 A&B**). For Ali-BC, FLowSOM and Louvain found a general immune cluster which showed a mix of CD45, CD3 and CD68 markers (**Fig. 3A**). For Jackson-BC, FlowSOM found two clusters that corresponded to T cells and general immune cells, while Louvain found a cluster which corresponded to B cells (**Fig. 3B**). Majority of clusters found by all methods represent luminal epithelial cells. In both datasets, hmiVAE identified a distinct basal epithelial cell cluster with clear expression of CK5 and CK14. Louvain clustering found two basal epithelial cell clusters in the Ali-BC one of which shows a mix of other stromal markers like SMA while the other only shows expression of CK14. In the Jackson-BC dataset, Louvain finds a basal epithelial cell cluster only showing expression of CK14 with low expression of CK5 (**Fig. 3**). For both datasets, FlowSOM resulted in many clusters which did not show high expression of any markers and therefore could not be assigned to any cell types represented by the marker panels.

**Figure 3.**
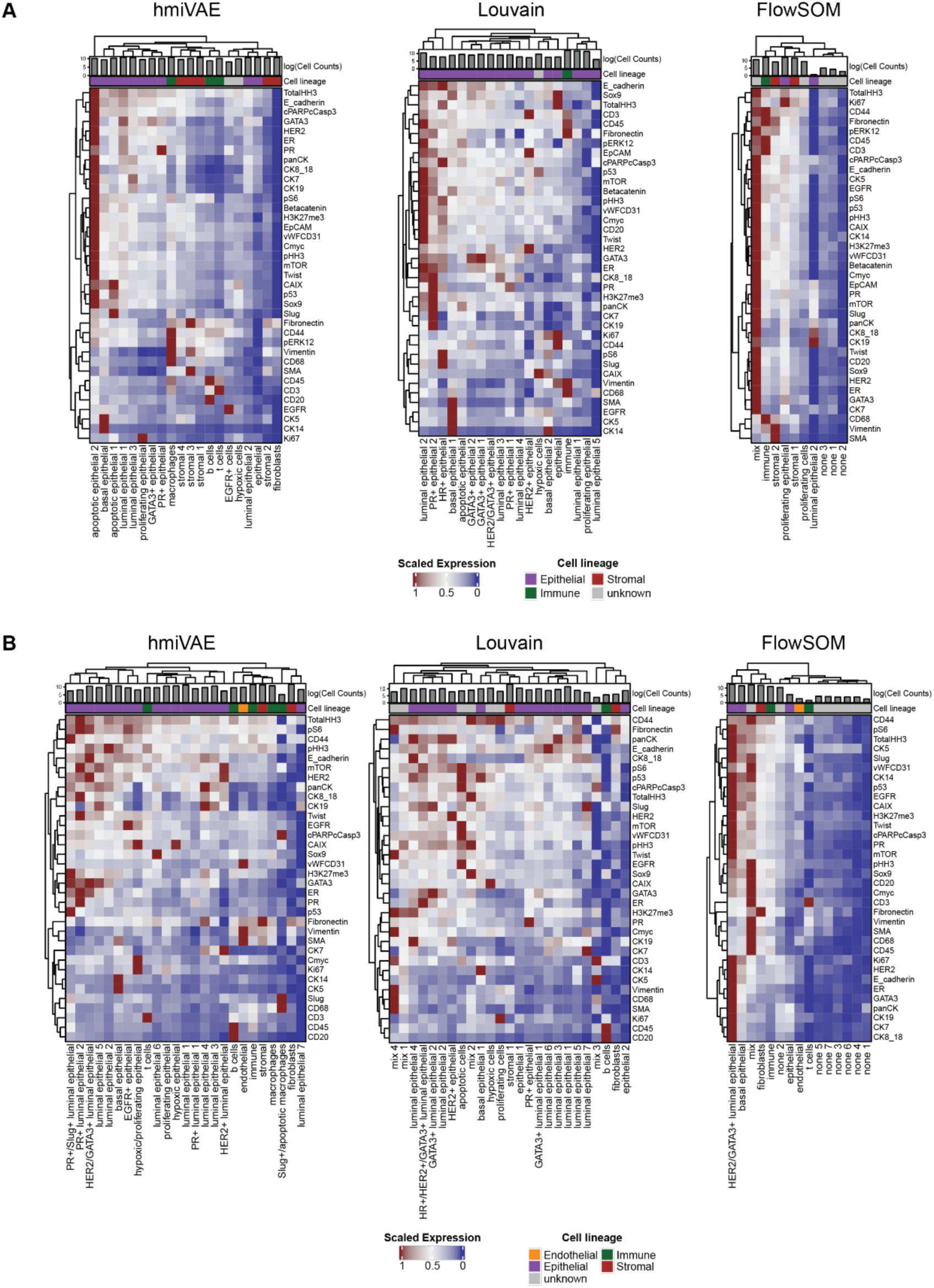
Cell type interpretation using expression-specific embeddings from hmiVAE and comparisons with Louvain and FlowSOM. Heatmaps showing expression values of expression-only features scaled to be between 0 and 1. Cell counts within each cluster are shown as logarithmized cell counts. **A** Cell types identified using hmiVAE vs identified by Louvain and FlowSOM in Ali-BC using only the expression features set. FlowSOM finds clusters with a mix of cell type markers, with one generalised immune cell cluster and some clusters that do not show high expression for any markers. Louvain also finds a general immune cell cluster and luminal and basal epithelial cell clusters. Clusters from the expression-only embeddings from hmiVAE finds specific immune cell clusters corresponding to B cells, T cells and macrophages, as well as luminal and epithelial cell clusters. **B** Cell types identified using hmiVAE vs those identified by Louvain and FlowSOM in Jackson-BC using only the expression features set. Similar to Ali-BC, FlowSOM finds clusters showing a mix of markers with two clusters corresponding to T cells and general immune cells, one cluster of endothelial cells and many clusters which do not show high expression for any markers. Louvain finds a cluster corresponding to B cells and clusters corresponding to basal and luminal epithelial cells. As before, clusters from the expression-only embeddings from hmiVAE finds clusters corresponding to B cells, T cells and macrophages, as well as clusters corresponding to endothelial cells, basal and luminal epithelial cells.

To further compare the clusters found by hmiVAE to those found by Louvain and FlowSOM, we treated the single-cell type assignments from Jackson et. al (2020)^5^ and Ali et al. (2020)^6^ as the ground truth labels for the Jackson-BC and Ali-BC datasets. For the Jackson-BC datasets, we also retrieved manual single-cell annotations by two independent annotators for a subset of 500 cells from Geuenich et al. (2021)^14^. We contrasted clusters with ground truth using the Adjusted Rand Index (ARI), a metric that quantifies the similarity between two clusterings of data points^38^. We found that for both datasets, hmiVAE achieves the highest ARI when compared to Louvain and FlowSOM indicating that it is better able to recapitulate the original, published cell type assignments (**Fig. 4C**). For the Ali-BC dataset, the ARI for hmiVAE is comparable to FlowSOM, but this is expected since FlowSOM was used in the original publication for clustering (**Fig. 4B**). In the comparisons with the manual-labelled single cell annotations, we found that only by using the expression embedding from hmiVAE to cluster, we were able to correctly identify B cells, macrophages and endothelial cells, while Louvain and FlowSOM failed to correctly identify these cell types in the subset of 500 cells. Louvain showed higher precision but similar recall in calling basal epithelial cells while hmiVAE had higher precision but lower recall for stromal cells. All methods had very similar precision and recall for luminal epithelial cell populations. This highlights that the expression-only cell representations learnt by hmiVAE may allow for the discernment of subtle immune and epithelial cell subtypes.

**Figure 4.**
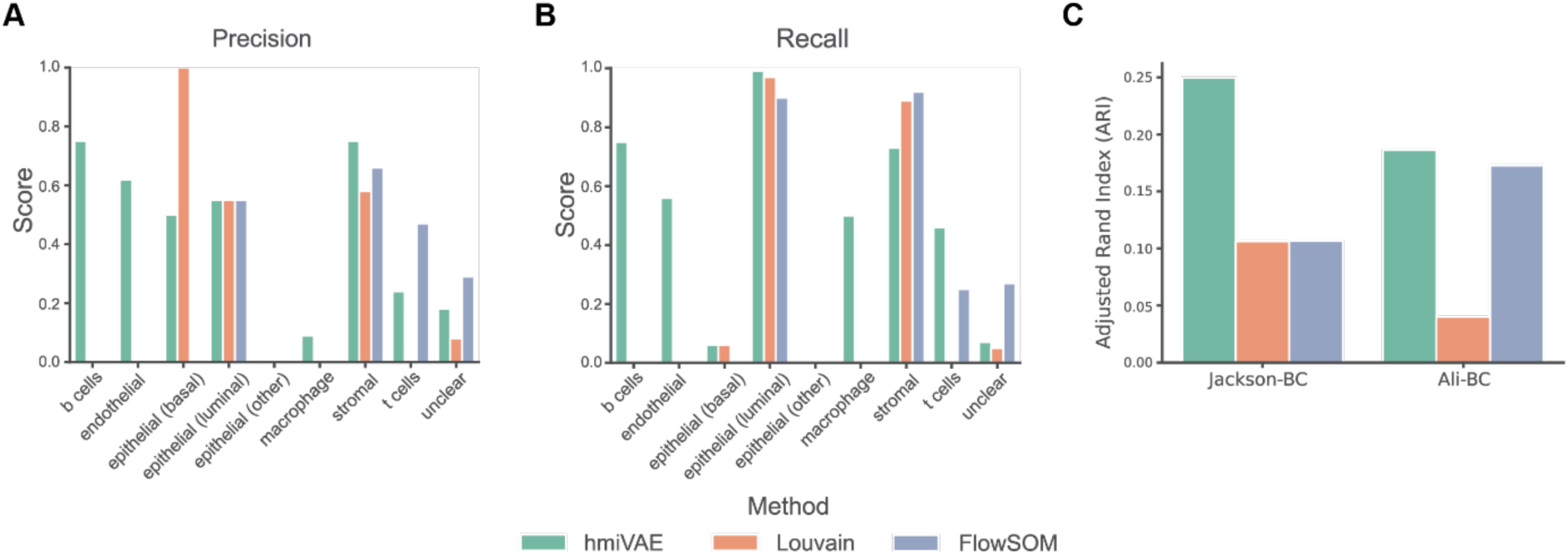
Comparisons of expression-only clusters found by hmiVAE, Louvain and FlowSOM. Using manual ground truth annotations for Jackson-BC, clusters found using the expression embedding of hmiVAE had higher precision in calling immune cells (B cells and macrophages), endothelial, and stomal cell populations **(A)** and higher recall for all immune cell populations in the dataset **(B)**. **C** Comparison to published annotations. When comparing to single-cell annotations from publications, hmiVAE was best in recapitulating the annotations for both datasets when compared to FlowSOM and Louvain.

### Clusters from integrated space are associated with clinical variables and patient survival

Existing approaches have used the latent space of a deep unsupervised models to encode spatial information and marker expression to derive patterns that provide clinically meaningful differences between patient groups^18,29,39,40^. Therefore, we examined whether the presence of the integrated clusters was associated with clinical variables such as stage, grade and cancer subtype (**Supplementary Table S1**) and survival. We first established two metrics quantifying the presence of cells from a specific cluster within a patient’s tissue sample: (i) cluster proportions, which can be described as the relative abundance of a cluster compared to others, and (ii) cluster prevalence per mm^2^ of tissue, which can be seen as a measure of a cluster’s density within the tissue (**Fig. 5A**, **Methods**). We calculated these metrics both the integrated clusters from hmiVAE as well as a baseline of the FlowSOM and Louvain clusters when using the full input feature set. We also compared with the clusters found by hmiVAE using only the expression feature embedding and the clusters found by FlowSOM and Louvain using the expression features only. For both prevalence measures, we computed its association with each cluster for each dataset. We also applied a Cox proportional hazards model to identify the clusters that were predictive of survival after stratifying on cancer stage (**Methods**).

**Figure 5.**
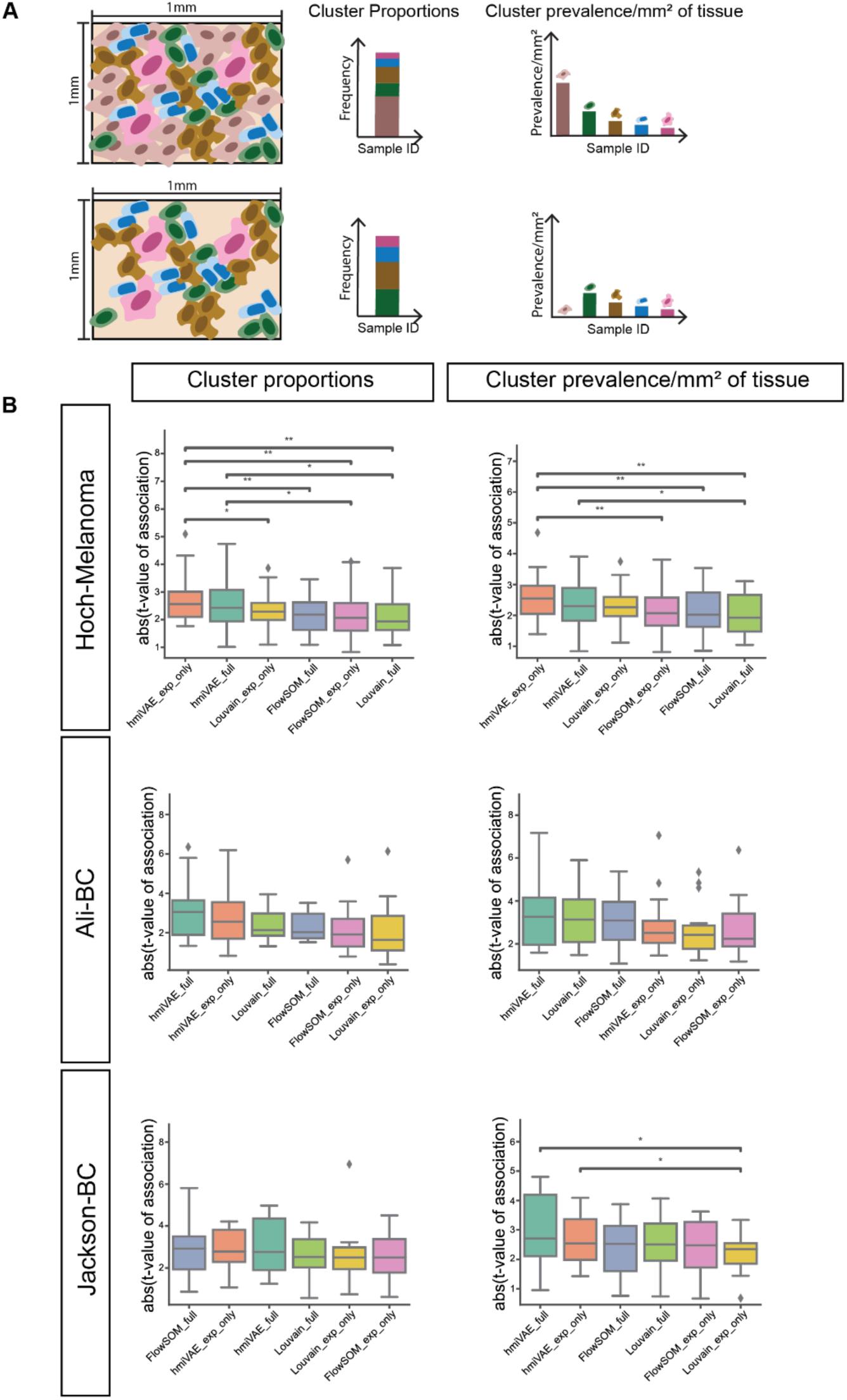
Clinical association with hmiVAE latent space. **A** Measures of cluster prevalence within a patient sample. We used two methods to describe cluster prevalence. (i) Cluster proportion - proportion of cells belonging to a given cluster within a patient and (ii) Cluster prevalence per mm^2^ of tissue or *tissue prevalence* - the abundance of cells from one cluster relative to the overall size of the tissue image. **B** Clinical variable associations across methods for all datasets. hmiVAE, using either clusters from the expression embedding or integrated latent space, found clusters with higher associations than Louvain and FlowSOM. Both Louvain and FlowSOM found clusters with higher clinical associations when run over the full features set. P-value annotation: *: 0.01 < p <= 0.05; **: 0.001 < p <= 0.01; ***: 0.0001 < p <= 0.001; ****: p <= 0.0001. All n.s. comparisons are omitted.

Across the Jackson-BC and Ali-BC datasets, we observed that although the difference between the mean cluster association with clinical variables across methods was not statistically significant, hmiVAE integrated clusters exhibited stronger associations with clinical variables compared to clusters from other methods, as assessed by both measures (**Fig. 5B**). For the Hoch-Melanoma dataset, we found the cluster proportions for the expression-only clusters had significantly higher associations with clinical variables than the clusters found by running FlowSOM and Louvain on only the expression (**Fig. 5B**). Our findings suggest that the integrated latent space learned by hmiVAE may provide a more robust representation of cellular phenotypes associated with clinical variables compared to traditional clustering methods.

To quantify the association between cluster prevalence and patient survival, we evaluated the concordance indices of Cox proportional hazards (CoxPH) models applied to clusters obtained as described above while controlling for disease stage. We observed that the hmiVAE integrated clusters consistently showed the highest concordance indices across all datasets, indicating better predictive performance in survival analysis (**Fig. 6A**). Furthermore, the superior performance of hmiVAE integrated clusters was statistically significant compared to directly clustering the entire feature set, particularly when using the cluster prevalence per mm^2^ of tissue. This significance was consistent across all datasets, suggesting that the integrated latent space learned by hmiVAE captures meaningful biological variation associated with patient outcomes. Specifically, in the Ali-BC dataset, the difference in concordance index between the hmiVAE integrated clusters and other methods was statistically significant using both measures, further supporting the robustness of our model.

**Figure 6.**
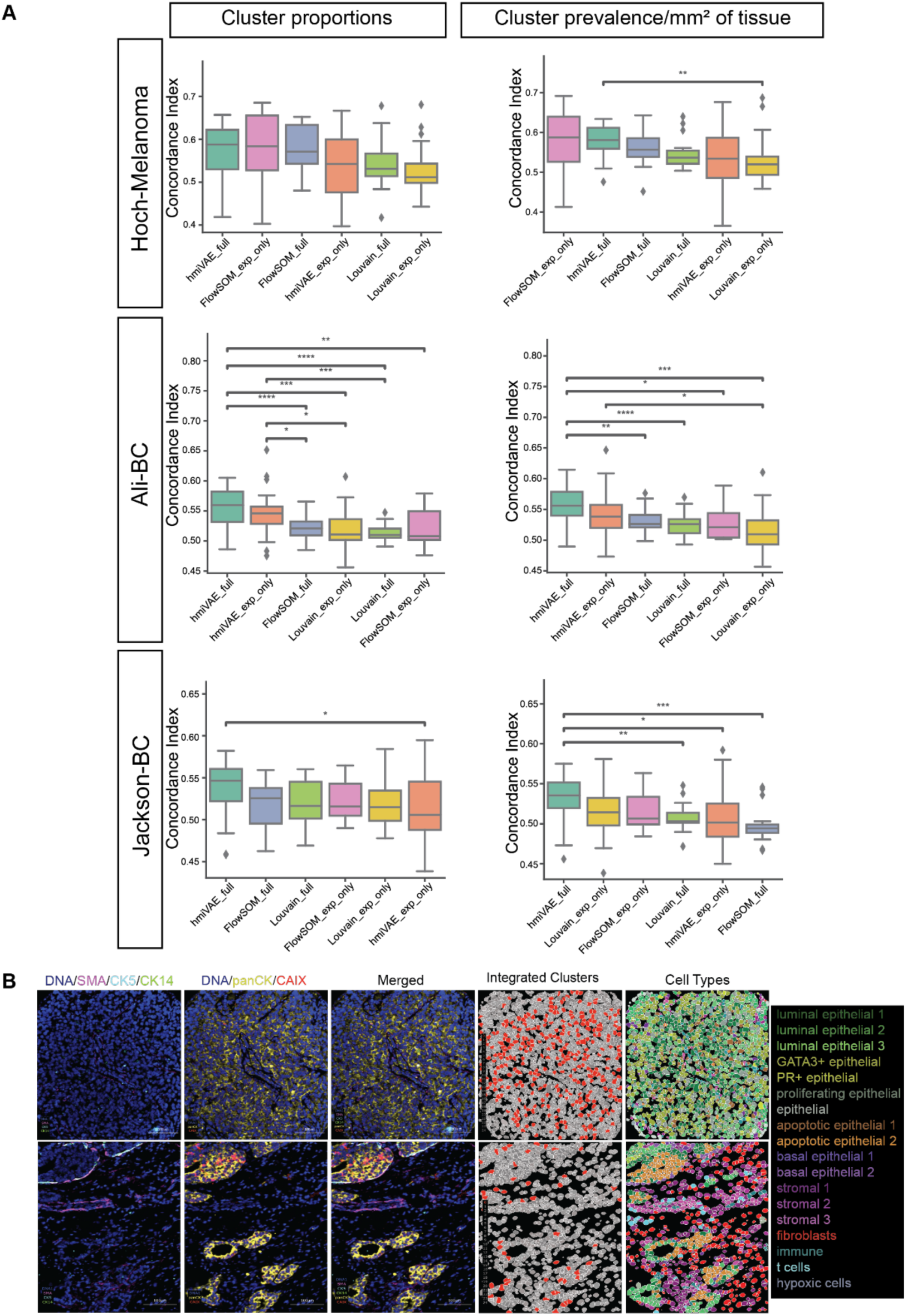
Survival analysis across methods for all datasets. **A** IMC images from the Ali-BC dataset. Images showing DNA, panCK, CAIX, SMA, CK5 and CK14 expression, as well as cell masks coloured by cell type and hmiVAE integrated space cluster ID for a patient with low number of cells belonging to integrated space cluster 0 **(bottom)** and a patient with high number of cells belonging to integrated space cluster 0 **(top)**. **B Concordance index for CoxPH models for each dataset across methods.** Clusters from the latent space learned by hmiVAE show higher concordance in comparison to Louvain and FlowSOM across all datasets, especially using cluster prevalence per mm^2^ of tissue to summarise cluster presence within a patient sample. P-value annotation: *: 0.01 < p <= 0.05; **: 0.001 < p <= 0.01; ***: 0.0001 < p <= 0.001; ****: p <= 0.0001. All n.s. comparisons are omitted.

We next interpreted the biological features of the clusters with significant hazards ratios for both cluster prevalence measures in Ali-BC and Hoch-Melanoma (**Supplementary Fig. 4&5**). In the Ali-BC dataset, hmiVAE found a cluster (*integrated cluster 0*) whose higher prevalence was significantly associated with higher risk based on a Cox proportional hazards model for both cluster proportion and tissue prevalence. According to marker expression, this cluster consists of luminal epithelial cells expressing the hypoxia marker, CAIX, and the proliferation marker, Ki67. Given the depletion of myoepithelial cells (expressing SMA, CK5 or CK14) in this cluster and its spatial context (**Fig. 1**), we hypothesized it may represent the hypoxic tumour core^41^. To investigate this further, we examined images from patients with high and low presence of cluster 0 in their tissue sample (**Fig. 6 & Supplementary Fig. 10**). We found that samples with a high presence of cluster 0 had high expression of the tumour cell marker panCK and hypoxia marker CAIX, with little to no expression of SMA, CK5 or CK14. Also, in the few regions that showed expression of SMA, the luminal epithelial cells were not surrounding the tumour cells. On the other hand, for patients with a low number of cluster 0 cells, samples had high expression of SMA, CK5 and CK14, low expression of panCK or CAIX, and the luminal epithelial cells were surrounded by myoepithelial cells (**Fig. 6B**). This spatial pattern supports our hypothesis that cells belonging to cluster 0 found by hmiVAE might belong to the tumour core.

In the Hoch-Melanoma dataset, *integrated cluster 10* was significantly associated with survival for both measures whereas the hazard ratio for *integrated cluster 15* was only significant using cluster proportions. We found that cluster 10 represented tumour cells expressing and surrounded by cells expressing IDO, which has been implicated in degrading T cell function^42^. These tumour cells were also surrounded by T regulatory cells (given by the neighbourhood expression of FOXP3) and macrophages (denoted by the neighbourhood expression of CD11b), both of which are canonical markers of a immunosuppressive microenvironment^43^. In contrast, cluster 15 represented tumour cells without such markers in their neighbourhood. A patient sample that had a high number of cells belonging to both clusters exhibited a mixed pattern with cells closer to IDO expressing cells belonging to cluster 10 and those away from these cells belonging to cluster 15 (**Supplementary Fig. 7**). We note that such spatially informed patterns would be difficult to identify using mean protein expression alone, which highlights the importance of incorporating neighbourhood information to identify cell states.

Our analysis demonstrates that the integrated clusters derived from the hmiVAE framework exhibit significant associations with various clinical variables and patient survival outcomes across multiple datasets, highlighting their potential as robust biomarkers for clinical stratification. Further investigations with larger datasets and additional statistical tests may provide deeper insights into the significance of these findings. Our results demonstrate the value of incorporating multi-view information and leveraging deep learning approaches for more accurate and clinically relevant insights into cellular phenotypes and their associations with patient outcomes in complex biological systems.

### Latent space projection with hmiVAE

The latent space of a VAE captures the salient features that explain the variation in high dimensional data^26,44^. Therefore, given shared original features in the high dimensional space, we sought to investigate how similar and transferable the learnt latent space and feature embeddings were between two IMC datasets. To do so we used the Ali-BC and Jackson-BC datasets, which share an antibody panel. We trained hmiVAE on one dataset (reference dataset) and used it to generate latent space and view-specific cell representations for the other dataset (query dataset). We did this by first clustering cells from the reference dataset using their latent space representations or view-specific embeddings and assigning them to a cluster ID. These latent cell representations or view-specific embeddings and their associated cluster IDs were then used to train a K Nearest Neighbour model (KNN)^45^, we call this the hmiVAE-KNN model. We then used the hmiVAE model trained on the reference dataset to generate latent space representations and view-specific embeddings for cells in the query dataset and these were then fed into the trained KNN model to generate cluster IDs. This approach was compared to a baseline model where we clustered the cells from the reference dataset using their original features— full (including features from all views), expression, nuclear co-localization, morphology and spatial context—and trained the KNN model using the resulting cluster IDs, we call this the baseline-KNN model. We then used this baseline-KNN model to generate cluster IDs for cells in the query dataset using their original features (**Methods**). For ground truth comparisons in the case of using the latent space learnt by hmiVAE, we compared the labels for the query dataset from the hmiVAE-KNN model to the cluster IDs resulting from training hmiVAE on the query dataset and clustering the cells to assign cluster IDs. In the baseline-KNN case, we used cluster IDs generated by directly clustering the cells from the query dataset using their original features for comparison. We applied KNN using both cosine and Euclidean distance metrics and compared clustering results to ground truth (hmiVAE or baseline) using the Adjusted rand index (ARI) and a balanced ARI score which adjusts the ARI score to weight underrepresented and overrepresented classes similarly^46^ (**Fig. 7A**).

**Figure 7.**
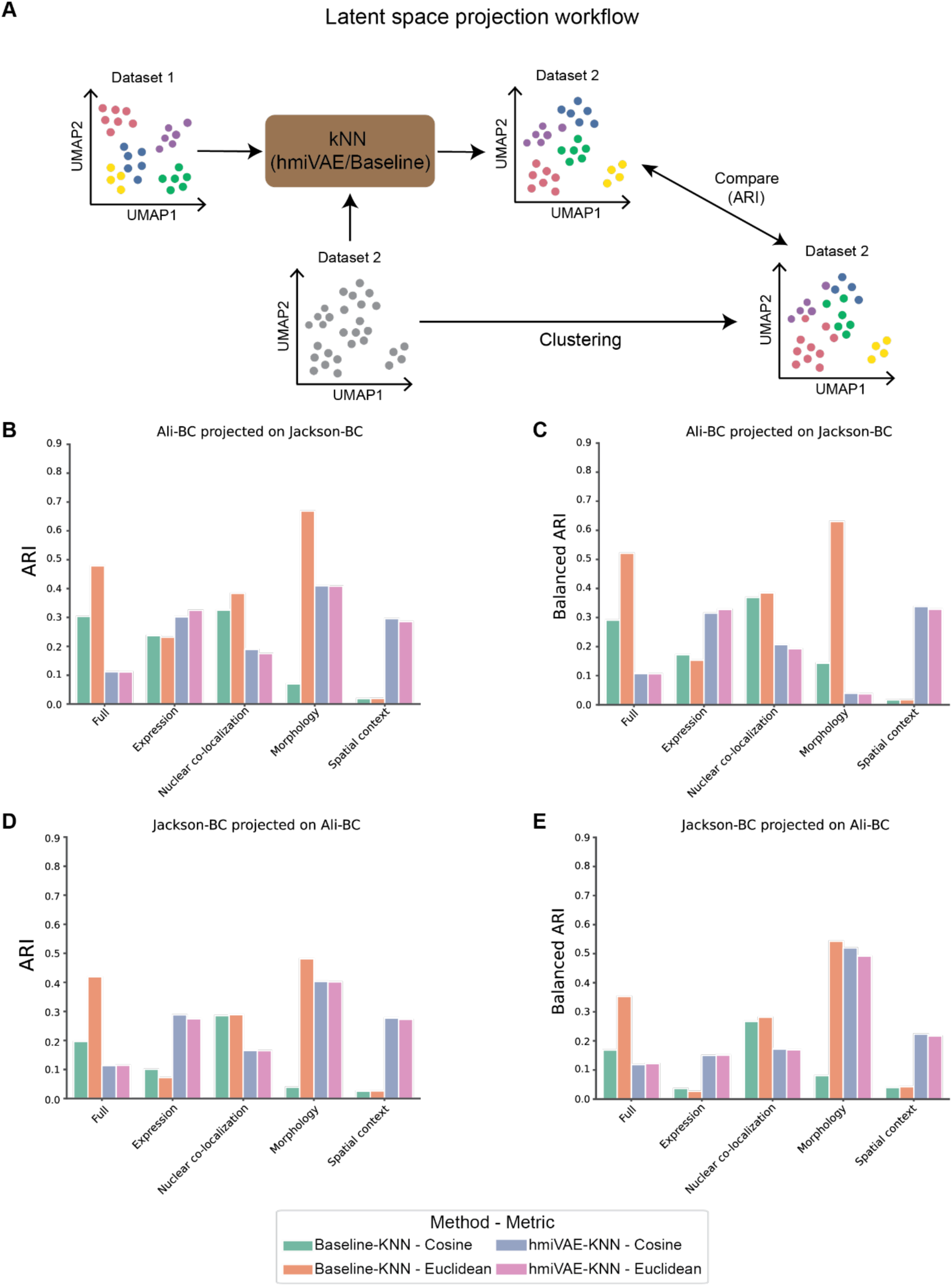
Latent space projection analysis with hmiVAE and comparisons with baseline KNN. **A** Schematic showing latent space workflow. A KNN model is trained using (i) labels from latent space or view-specific embeddings of hmiVAE trained on Dataset 1 (hmiVAE-KNN) or (ii) labels from clustering directly on the full or view-specific features set from Dataset 1 (baseline-KNN). Labels for an independent dataset, Dataset 2, are generated from this trained KNN model. The KNN clustering results are compared to cluster labels from ground truth Dataset 2 using adjusted rand index (ARI) or balanced ARI. **B - C** Projection results for Jackson-BC as Dataset 1 and Ali-BC as Dataset 2 using both cosine and euclidean distance metrics for KNN reported as ARI **(B)** and balanced ARI **(C). D - E** Projection results for Ali-BC as Dataset 1 and Jackson-BC as Dataset 2 using both cosine and euclidean distance metrics for KNN reported as ARI **(D)** and balanced ARI **(E)**. Full - integrated space or full features set.

Our analysis revealed distinct performance trends based on the feature sets used. ARI and balanced ARI also did not differ much except in the case of morphology-only features for hmiVAE-KNN especially when projecting cells from Ali-BC onto the morphology-only embedding learnt from Jackson-BC (**Fig. 7B&C**). We notice that the baseline-KNN model consistently achieved higher ARI and balanced ARI scores when applied to the full feature set and nuclear co-localization features in both datasets. While hmiVAE exhibited superior performance in clustering when leveraging expression and spatial context features (**Fig. 7**). Furthermore, the choice of distance metrics did not influence the projection capability of the features learnt by hmiVAE but did when using the original features, especially for the full feature set and morphology features. This delineation underscores the importance of considering which features are being used for projection. Latent space projection is considered a beneficial potential extension for applications in biology^47^, but our study demonstrates that this might not be so straightforward when projecting cells from a different dataset onto a latent space or view-/modality-specific embedding.

## Discussion

Here, we have investigated the use of VAEs for multi-view modelling of IMC data. Our approach revealed integrated clusters corresponding to diverse cellular phenotypes characterised by subtle variations in protein expression, subcellular localization, and microenvironmental context. Notably, our findings highlight the necessity of incorporating nuclear co-localization, morphological features, and spatial information to capture the complexity of cellular states more accurately than from using expression alone. We show that the view-specific embeddings and integrated latent representations for each cell led to the identification of clusters which had strong associations with clinical variables and patient survival, as well as improved performance for common downstream tasks such as cell type annotation. We note that while this investigation followed the standard approach of extracting single cells and identifying cell-level phenotypes, other studies have improved the performance on downstream tasks by creating pixel-level annotations and thereby increasing robustness of the features extracted^48^. This approach could be used to adapt our input features, such as pixel-level protein expression and spatial context, and to investigate the effects on the learnt representations and the performance on downstream tasks. Further, in this study, our exploration of hyperparameters in the hmiVAE architecture unveiled effects on reconstruction performance, emphasising the importance of dynamically adjusting the weight of the KL-Divergence term and optimising hidden layer size and batch size for superior reconstruction. Leveraging these insights, we identified optimal hyperparameter configurations for each dataset, maximising reconstruction likelihood to construct integrated clusters representative of the underlying cellular heterogeneity. However, we considered only a subset of possible hyperparameters that could be expanded in future work to contain the learning rate known to have a big impact on model performance^49^. In our case, it could also be beneficial to vary the size of the view-specific embeddings given the differences in the number of features for each input view.

In our comparisons, all three methods, hmiVAE, FlowSOM and Louvain tended to find clusters with higher associations with clinical variables when run on the set of features from all four views. Further, hmiVAE was able to find distinct spatial patterns with poor survival outcomes without any prior knowledge, indicating that this information is contained in IMC data and can be learned with deep learning approaches in an integrated way. This shows the importance of incorporating views beyond per-cell mean protein expression to learn cell states associated with disease. We expect that spatial patterns and novel cell states learned by hmiVAE can be related to transcriptome or genome data where there exist match patient samples. Such a study could reveal whether different genomic programs are present in patients exhibiting higher or lower prevalence of specific cell states or spatial patterns, thereby increasing our understanding of patient heterogeneity in disease.

Our analysis demonstrated that while hmiVAE performed well on integrating the different views of HMI data and a variety of downstream analysis tasks, the applicability of such a method was not without its limitations. The latent projection analysis showed that the learnt expression and spatial context embedding generalised well over datasets with similar panel designs, but this was not the case for the integrated latent space and the nuclear co-localization embedding, underlying the need to investigate batch effects during projection for the different views. It may also be beneficial to understand how network architecture affects this projection of new cells onto the latent space or view-specific embeddings. Further work is also required to determine how well the learnt latent space generalises over datasets with different panel designs, tissue types and disease conditions as well as which dataset characteristics may impact generalisation. Further, using a KNN model for projection analysis, especially of the latent space may not be suitable as it relies on distance metrics such as Euclidean distance or cosine similarity to determine closeness of points. These metrics for similarity have been seen to lead to arbitrary results when applied to learned latent space vectors^50^. It is worth noting that finding effective quantitative benchmarks and metrics for deep learning methods using HMI data is challenging since there is a lack of ground truth annotations as it involves time and expertise which might not be readily available^51^. Further, established evaluation metrics do not exist for assessing the biological validity of the found clusters and so some studies create their own metrics^22^, but these may be hard to replicate in different studies. Therefore, there is a need to create robust and reliable metrics and labels for testing and validating the application of deep learning methods to HMI data, which will give us a better understanding of their utility beyond manual inspection and visualisation.

Overall, our investigation shows that deep learning methods such as VAEs can successfully integrate different views of HMI data to learn cell state representations associated with clinically relevant variables in cancer. Deep learning methods for HMI data are still in their infancy and our work highlights their potential to improve understanding of patient-level disease presentation by leveraging rich information captured by HMI technologies.

## Methods

### Datasets used

The Jackson-BC dataset comprised 358 images obtained from a Tissue Microarray (TMA) of tissue sections sourced from breast cancer patients, encompassing 802,591 cells and incorporating 36 proteins, including two DNA intercalators^5^. The Ali-BC dataset, derived from the METABRIC study^52^, featured 548 images from a TMA of tissue sections of patients with primary invasive carcinoma. This dataset included 536,883 cells and utilised 39 stains, including two DNA intercalators^6^. Notably, both datasets shared an antibody panel designed to target epitopes specific to breast cancer, encompassing markers for cell cycle regulation, phosphorylation-based signalling, and distinctive markers for epithelial, endothelial, mesenchymal, and immune cell types. The Hoch-Melanoma dataset comprised 167 images extracted from a TMA of biopsies originating from patients with stage 4 or stage 3 melanoma. This dataset encompassed 989,404 cells and employed 46 stains, incorporating two DNA intercalators. The antibody panel for the Hoch-Melanoma dataset included markers for tumour cells, alongside those targeting various immune cell types and their activation states^31^. Each dataset was accompanied by comprehensive clinical and survival information, detailed in table S1.

### Feature derivation

Each dataset consists of a: (1) 3-dimensional expression array which contains the pixel-level protein expression counts, (2) cell segmentation mask array which contains cell ID assignments for each pixel and (3) channel-to-protein identification dataframe which associates each z-stack in the expression array to a protein measured in the experiment. The mask and expression arrays have the same x and y coordinates, and the size of the z-axis in the expression array corresponds to the number of channels in the channel file.

Expression features: To create the protein mean expression features, we subset to pixels belonging to the same cell ID (as contained in the mask array) in the expression array for each protein. We average over the counts over these pixels for each protein to give the average count for expression within that individual cell. We do this for each cell creating a *N* x *P* matrix where *N* is the number of cells in the dataset and *P* is the number of proteins measured in the experiment. We denote this matrix **Y**.

Protein nuclear localization score features: Due to the resolution of IMC, we restricted protein subcellular localization to in-nucleus or not in-nucleus. Therefore, this should be treated as a protein nuclear localization score. We generate this score by first creating a “mean nuclear stain” by taking the average of the two Iridium DNA stains for each pixel. We subset to pixels belonging to the same cell ID and then do a pixel-wise correlation between each protein and this mean nuclear stain. A high correlation means that the same pixels within a cell have high expression of a given protein and a high mean nuclear stain, suggesting that this protein is expressed in the nucleus. We do this for each cell creating a *N* x *P* matrix where *P* is the number of correlations (note here that the number of correlations equals the number of proteins measured in the experiment). We call this matrix **S**.

Morphology features: We use Python’s regionprops function from the scikit-image library^53^ to create the morphology features: area, perimeter and eccentricity. We also adopt the concavity and asymmetry features from DeepCell^11^. We compute these features for each cell, thereby creating a *N* x *M* matrix where M is the number of morphology features. In our case, *M* = 5. We call this matrix **M**.

Spatial context features: For the spatial context features, we first compute the 10 nearest neighbours based on spatial coordinates for each cell using scanpy^35,45^. Using these neighbours, we create a *N* x *N* sparse matrix **D** s.t. **D***_ij_* = 1 if cell *i* is the neighbour of cell *j* and 0 otherwise. We then scale **Y**, **S**, and **M** to be such that each feature in these matrices has a mean 0 and variance 1 and concatenate these together. The sparse matrix **D** is then multiplied with this concatenated matrix, with each component normalised by the number of each cell’s neighbours. This gives us a *N* x *L* matrix, where *L* = *P* + *C* + *M*. We call each row the spatial context of a cell, and denote this matrix by **C**.

Background features and one-hot encoding: We create a sample-wise one-hot encoding for each cell i.e., for a given cell, the vector is of length *R* for *R* number of samples and has a 1 in position *r* if the cell belongs to sample *r* and 0 otherwise. Along with the one-hot encoding, we create several background covariates. (1) When background stains such as ArAr80 were available, we computed the mean background channel staining per cell and correlated this mean background stain with the mean protein expression per cell for all proteins. A high correlation between the background stain and a protein’s expression across cells would suggest a non-specific staining for that protein. (2) Since staining efficiency can vary between tissue samples^32^, we also computed the ‘mean sample intensity’ which we define as the average pixel intensity over all channels for an image. (3) We also computed the average value of signal intensity across all the background channels per sample.

In the case where information about background stains was not available, we computed the sample intensity and the average background stain of each protein. These background stains were defined as the mean over a protein’s expression values for 0-labelled pixels (in the mask file, pixels belonging to a cell are denoted by a non-zero integer corresponding to that cell’s ID, whereas pixels that are labelled as “0” are treated as background). We denote all background features and one-hot encoding covariates by *b*.

### hmiVAE Architecture

hmiVAE follows a Variational Autoencoder architecture^26^. It consists of two neural networks: an encoder and a decoder. The encoder takes in a data matrix corresponding to each view, along with the additional covariates outlined in *Background features and one-hot encoding* and generates a separate embedding for each of the four views: protein mean expression, protein subcellular localization, cell morphology and spatial context. These view-specific embeddings are then concatenated, passed through an additional hidden layer and integrated to create a latent representation *z* for each cell informed by all four views. The decoder takes in the latent representation of each cell and any additional covariates input to the encoder, i.e. the input to the decoder is concatenated as *z + b*. The non-linear decoder follows a symmetric architecture to the encoder in which integrated hidden layers are followed by view-specific hidden layers that reconstruct each view separately.

### Model training

Hyperparameter tuning: We tuned the model by varying 6 hyperparameters: (i) initialization (random seed – 0, 1, 42, 123, 1234), (ii) number of hidden layers (1 or 2), (iii) hidden layer size (i.e. number of nodes in each hidden layer – 8, 32, 64), (iv) latent representation dimension (10 or 20), (v) beta scheme (i.e. warmup of the weight of the KL-term in the Evidence Lower Bound (ELBO) loss, starting at zero and incrementing by 0.1 after each epoch, or keeping the weight of the KL-term constant at 1.0) and (vi) batch size (i.e. 40000, 16000, 8000, 5333, 4000). We optimized the ELBO loss function to train our method as defined below:

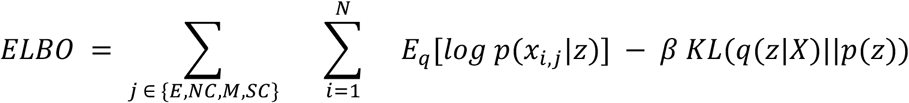

In the above equation, the left-hand side of the expression is the reconstruction likelihood for all views summed, the right-hand side is the KL-divergence between the estimated distribution, q, and the true distribution, p. The feature vector for each view j and cell i is represented by x_i,j_, z is the latent space feature vector and X is the set of all feature vectors or data i.e. *X* = {*x_E_*, *x_NC_*, *x_M_*, *x_SC_*}.

We created train and test splits for each dataset and used a grid search algorithm over the different combinations of these hyperparameters and selected the combination that resulted in the best reconstruction likelihood over test data.

Run with best model: hmiVAE was run on each dataset with the combination of hyperparameters that resulted in the best test likelihood for that dataset. The optimal combinations for hyperparameters for each dataset are detailed in **Supplementary Table S2**. The whole dataset is passed through the trained model to generate the latent representations as well as the view-specific embeddings for each cell in the dataset.

Clustering: For creating clusters, first we computed distances between all cells in the integrated and view-specific latent spaces and found the 100 nearest neighbours within the corresponding latent space for each cell using the neighbours function from scanpy^35^. Next, we use the leiden algorithm with resolutions 1.0, 0.5, 1.0, 0.1 and 0.5 for clustering over the latent space, expression, correlation, morphology and spatial context embeddings respectively. For the Ali-BC dataset, we clustered over the expression-only cell embedding using a leiden resolution of 1.0.

Benchmarking with FlowSOM and Louvain: Louvain is a community detection algorithm^36^ very similar to leiden^54^ and is popularly used in single-cell data analysis. To mimic the standard workflow in IMC cluster analysis, we first found the 10 closest neighbours for each cell over the full and expression-only feature space. We then tried different values for the resolution parameter in louvain to result in a number of clusters which was at most 1 away from the number of hmiVAE leiden clusters for easy comparison. For the Ali-BC dataset, a resolution of 0.3 for clustering over the expression-only feature space was used, for the Jackson-BC dataset a resolution of 0.14 was used and for the Hoch-Melanoma dataset, we used a resolution of 0.16. For the full feature set we used a resolution of 0.9 for Ali-BC, 0.4 for Jackson-BC and 0.15 for Hoch-Melanoma.

FlowSOM, on the other hand, uses self-organising maps to sort single cells into clusters^37^ and requires users to set the number of expected clusters. It does so iteratively until the cell membership of the clusters stop changing or max iterations are reached. We set the number of neurons for clustering over the full feature set and expression features to be equal to the number of hmiVAE clusters that were found using the integrated space and expression-only embedding respectively. We noticed that as not all neurons get cells assigned to them, FlowSOM ‘skips’ some cluster IDs and therefore, results in fewer clusters than hmiVAE or Louvain.

### Ranking of feature drivers per cluster and cell type annotation

Ranking features for integrated clusters: To identify the features driving the clustering in the integrated latent space of hmiVAE, we applied the rank_gene_groups function from scanpy^35^ using the features from each view separately. For each cluster, we selected the top three features which had a positive t-value and an adjusted p-value of less than 0.05. We combined all the features across all clusters and as this list would contain lots of repeated features, we subset the list to only unique features across the clusters and then selected their values for each cluster. For the final list of feature drivers across all clusters, we relaxed our criteria for statistical significance and t-value to include the values for those features which were not significantly different for some clusters among the list of unique features.

Annotation of cell types: For our cell type annotation we used the rank_gene_groups function from scanpy^35^ and used violin plots for visual confirmation after applying a StandardScaler scaling of the expression features. For the Ali-BC and Jackson-BC datasets, within the epithelial cell lineage, we used CK8/18, CK19, CK7 and panCK as markers for luminal epithelial cells and CK5 and CK14 for basal epithelial cells. CD31/vWF was used as a marker for endothelial cells, and Fibronectin was used to determine fibroblasts and general stromal cells along with Vimentin and SMA. For immune cell lineage, we used CD45 and CD20 for B cells, CD45 and CD3 for T cells and CD45 and CD68 for macrophages. Some cell state markers were also included e.g. Ki67 for proliferation and CAIX for hypoxia.

For the Hoch-Melanoma dataset, the panel consisted of tumour cell markers (e.g. MiTF, SOX10, *β*-Catenin) and immune cell markers (e.g. CD3, CD8, CD303 etc.) and so, the cell types correspond largely to these lineages. The panel also consisted of markers for T cell activation such as CD45RO for memory T cells and CD45RA for naive T cells.

Any clusters that did not show any significant expression for the lineage markers as described above were annotated as “unknown”. We carried out the same workflow to annotate cell types identified by FlowSOM and Louvain by clustering over the expression-only features.

### Benchmarking cell type annotations

Comparisons using published dataset annotations: For the Ali-BC and Jackson-BC datasets, the single-cell assignments were retrieved from the relevant publications. For the Ali-BC dataset, we converted our annotations from hmiVAE, Louvain and FlowSOM into pandas dataframes, and merged it with the dataframe containing the Ali-BC publication labels using the sample ID and cell ID. Similarly, for the Jackson-BC dataset, we first matched each single-cell to its published meta cluster ID and created sequential cell IDs for our cell labels to match those from the publication as the labels were not sequential in the original masks which generated the cell IDs, however, it was a simple one-to-one matching to create the sequential labels. The dataframes containing the published cell labels and our annotations from hmiVAE, Louvain and FlowSOM were then merged as described earlier. The labels were compared using the Adjusted Rand Index function available from scikit-learn package^55^.

Comparisons using manual ground truth annotation of 500 cells: This was only performed for the Jackson-BC dataset as this was the only dataset that had any manually labelled ground truth annotations. These annotations were made by two independent annotators for a subset of 500 cells, further details on how the annotations were done are described elsewhere^14^. We edited the format of our cell IDs and merged the dataframes as described earlier and computed the precision, recall, specificity and F1-score for the cell type classifications over these 500 cells using the classification report and confusion matrix functions from the scikit-learn package^55^.

### Association with clinical variables

We did so by using two methods to define *cluster prevalence*: 1) We summarised the proportion of cells belonging to a sample from each cluster ID and 2) the prevalence of cells from each cluster per mm^2^ of patient tissue. We did this analysis for cluster IDs which resulted from clustering over the full feature set and only the expression features set for FlowSOM and Louvain. For hmiVAE we used the cluster IDs which resulted from clustering over the integrated latent space and the expression-only cell representations.

To get the proportion of cells belonging to a cluster ID, for each sample ID, we counted the number of cells that belonged to each cluster ID and divided that number by the total number of cells within a sample. This resulted in a number between 0 and 1 for each cluster ID and these numbers all summed up to 1. We call this *cluster proportion*.

To get the prevalence of each cluster as an instance per mm^2^, for each sample ID, we counted the number of cells that belonged to each cluster ID and divided it by the dimensions of the image belonging to the sample ID. To make computation easier, we multiplied the resulting number with 1e6 (assuming an image of size 1000×1000 pixels). We call this *cluster instances per mm^2^ of tissue*.

Association with clinical variables: We carried out an association study between the integrated latent space and the clinical variables for the Jackson-BC, Ali-BC and Hoch-Melanoma datasets. This was only done for hmiVAE as FlowSOM and Louvain only generate labels for each cell and do not generate any embeddings. To understand and compare the association between the resulting clusters, we conducted an association analysis between cluster prevalence and clinical variables for all three methods (hmiVAE, Louvain, FlowSOM) across all three datasets.

Association of latent space with clinical variables: For association between the latent space and clinical variables, for each sample in the dataset, we first calculated the median cell embedding for each dimension of the latent space. Next, we selected patients belonging to the same category for each clinical variable and ran a logistic regression with all the median latent space values to find the associations of each latent dimension with each category for a given clinical variable.

During this analysis, we found that the inclusion of some latent dimensions for a few clinical variables resulted in a singular matrix during computation, so we excluded them and re-ran a logistic regression with the remaining latent dimensions. For some categories of clinical variables, there were not enough samples so these were removed from the analysis.

Association of cluster proportion with clinical variables: Based on our definition of cluster proportion; in order to prevent a degenerate matrix in our analysis, we ran a logistic regression one cluster ID at a time using a 1 vs rest approach. We did this for all cluster IDs and all categories present within all clinical variables. Categories that did not have enough samples, we removed or recorded with a ‘NA’ t-value.

Association of cluster prevalence as instances per mm^2^ of tissue: For the association between cluster instances per mm^2^ of tissue, we did not have the same issue of a degenerate matrix as in the case of cluster proportion and found that the scaling by 1e6 aided in this. Therefore, we ran a logistic regression over all the cluster IDs for all categories within every clinical variable. Similarly, as before, categories that did not have enough samples were removed or recorded with a ‘NA’ t-value.

### Survival analysis using Cox proportional hazards models

To find the relationship between patient survival outcome and cluster prevalence, we used the python lifelines package^56^ to fit a Cox proportional hazards model including disease stage as a covariate. We carried out this analysis for each cluster ID at a time, for both cluster proportions and cluster instances per mm^2^ of tissue. *P* values for hazard ratios were adjusted by Benjamini-Hochberg correction.

### Projection on latent space and comparison with baseline

Only the Jackson-BC and Ali-BC datasets were included for this analysis as these share an antibody panel. We did this analysis in two ways: use the latent space learnt from the cells in Jackson-BC to generate low-dimensional cellular representations for cells in Ali-BC and vice versa. We will call the dataset that the latent space belonged to as the ‘reference dataset’ and the dataset that is projected as the ‘query dataset’.

While carrying out this analysis, we needed to be mindful of the different numbers of background stains and samples present in both datasets as this would change the number of covariates expected by the model. In the case where the query dataset had more background stains than the reference dataset, we selected the stains that matched between the two. While in the case where the reference dataset had more background stains than the query dataset, we selected all the stains present in the query dataset while setting the value for all ‘missing’ stains to zero. In both scenarios, we used a random one-hot encoding for samples similar to Gayoso et al. (2021)^22^. This ensured that the number of total covariates input to the model was the same between the reference and query datasets. Details of how this analysis was carried out is outlined in the *Latent space projection with hmiVAE* section of the main text.

## Data and code availability

Code required to reproduce this study can be found at https://github.com/camlab-bioml/hmiVAE_manuscript and code required to run the hmiVAE model can be found at https://github.com/camlab-bioml/hmiVAE.

## Contributions

Project conceptualization: SA, KRC, HJ. Data analysis and model development: SA, AS. Manuscript preparation: SA, KRC.

## Supplementary Materials

**Supplementary Figure 1.**
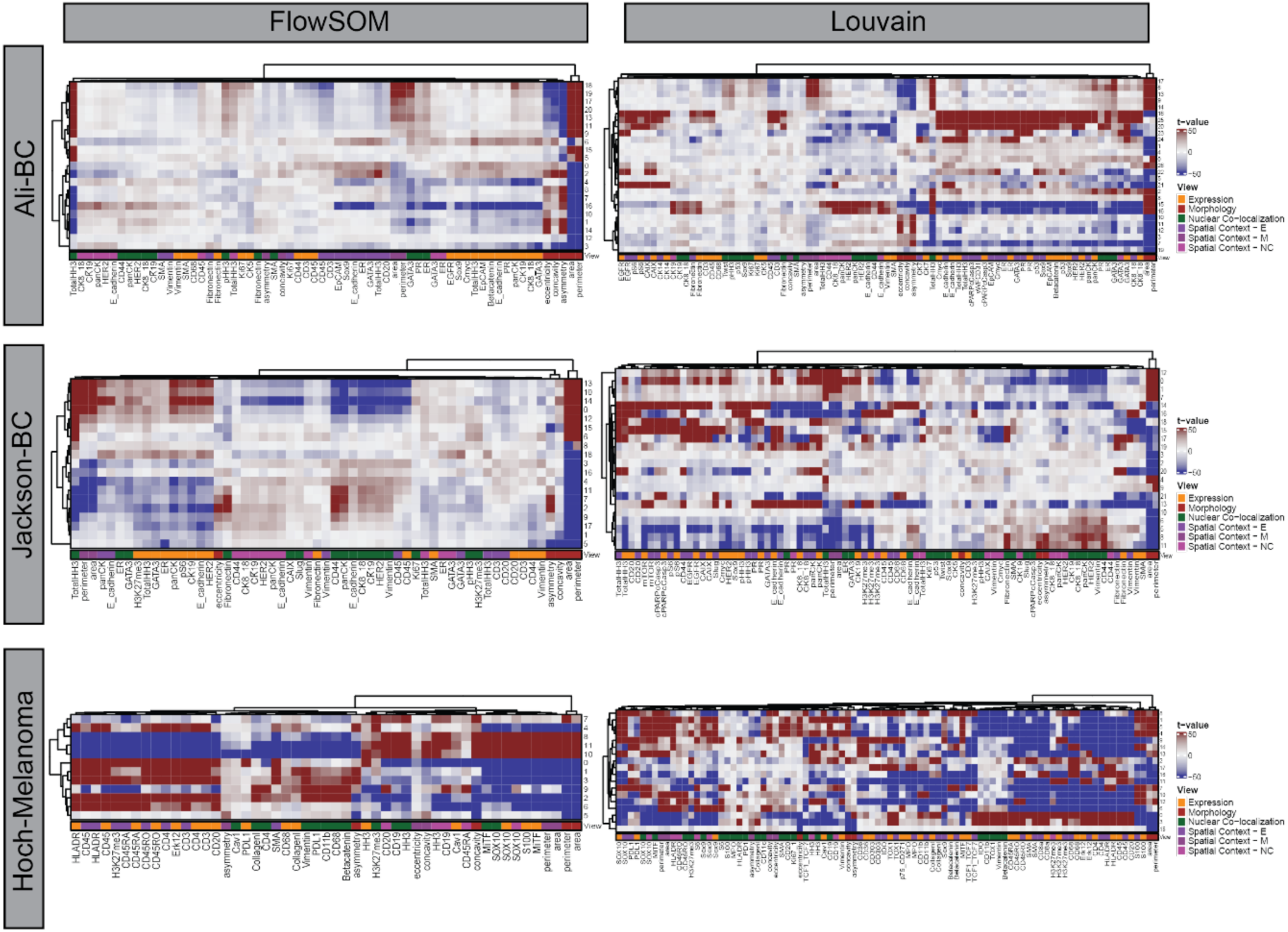
Ranking of all features for Louvain and FlowSOM across datasets.

**Supplementary Figure 2.**
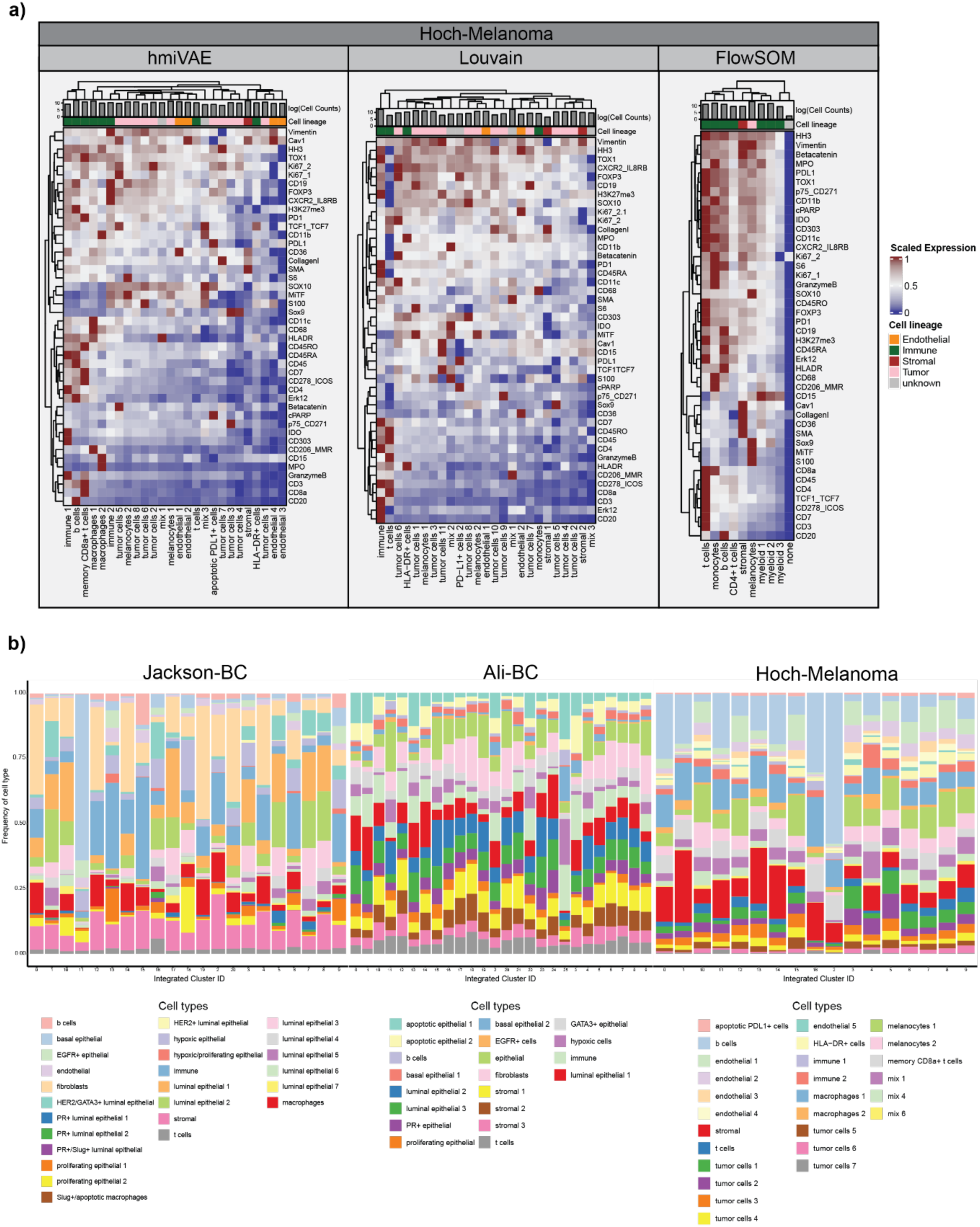
**A)** Cell types identified using hmiVAE vs identified by Louvain and FlowSOM in Hoch-Melanoma. **B)** Proportion of cell types occurring in each integrated cluster from hmiVAE.

**Supplementary Figure 3.**
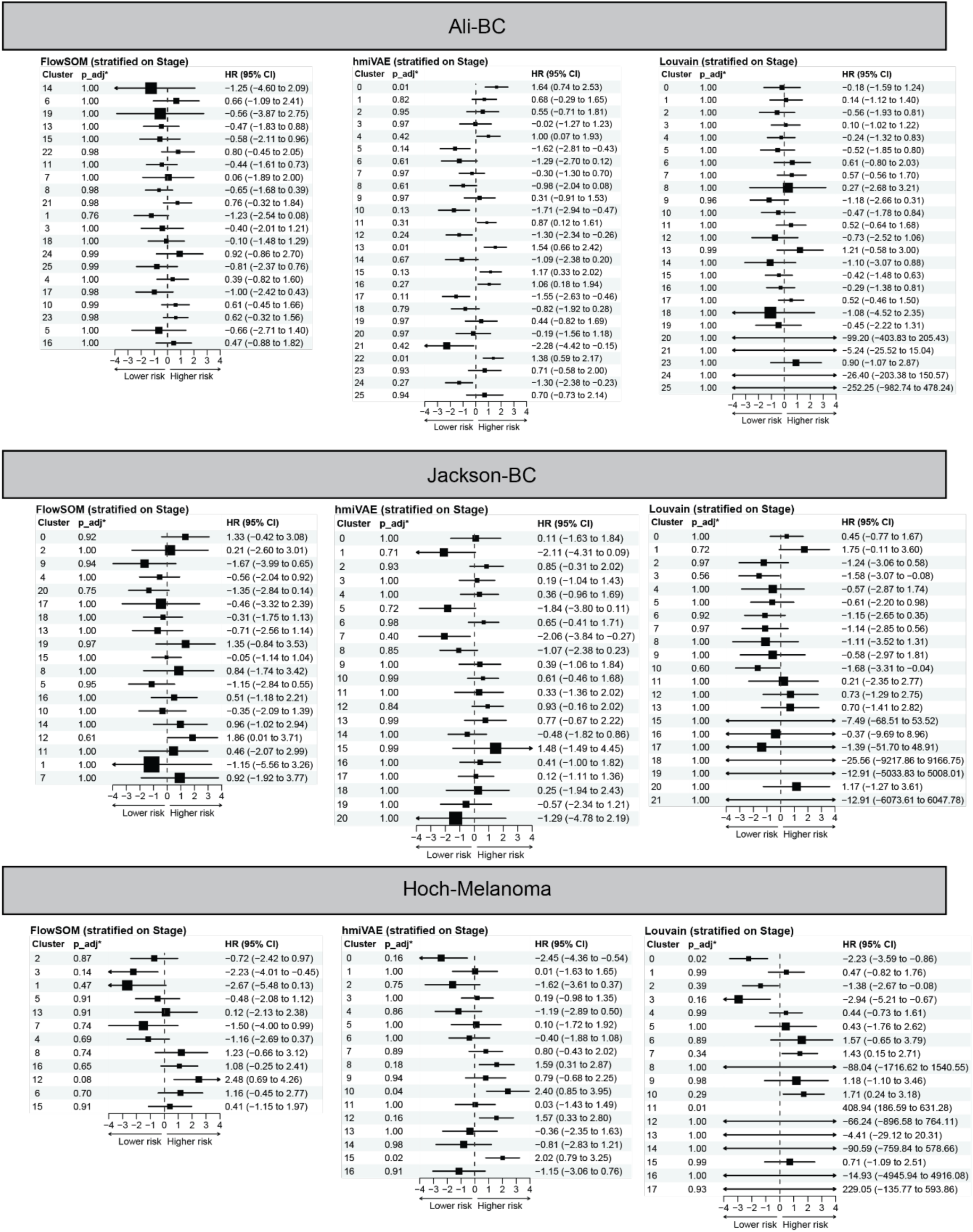
CoxPH model hazard ratios with cluster proportions for clusters from full feature set for Louvain and FlowSOM and hmiVAE integrated clusters for all datasets.

**Supplementary Figure 4.**
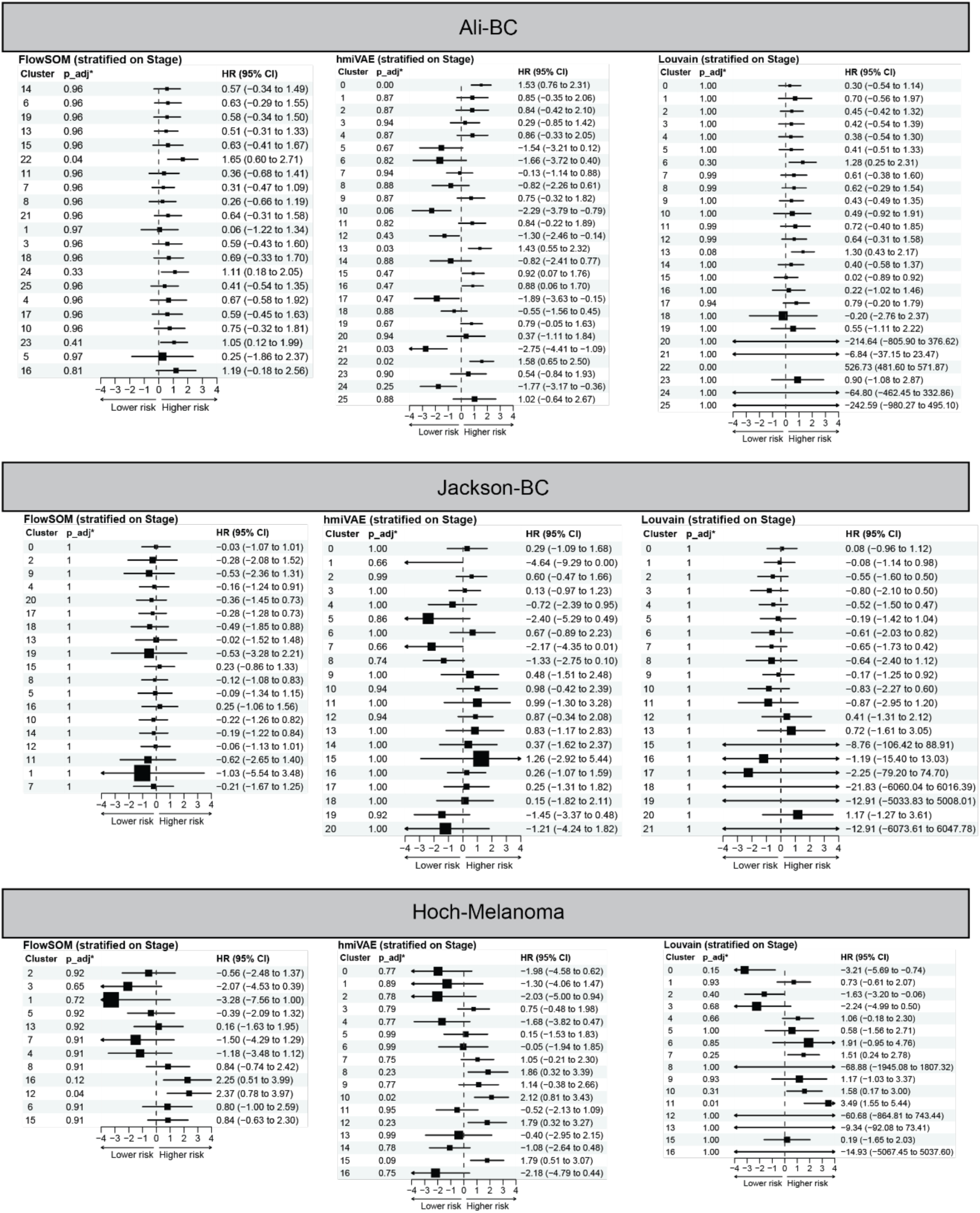
CoxPH model hazard ratios with cluster prevalence per mm^2^ of tissue for clusters from full feature set for Louvain and FlowSOM and hmiVAE integrated clusters for all datasets.

**Supplementary Figure 5.**
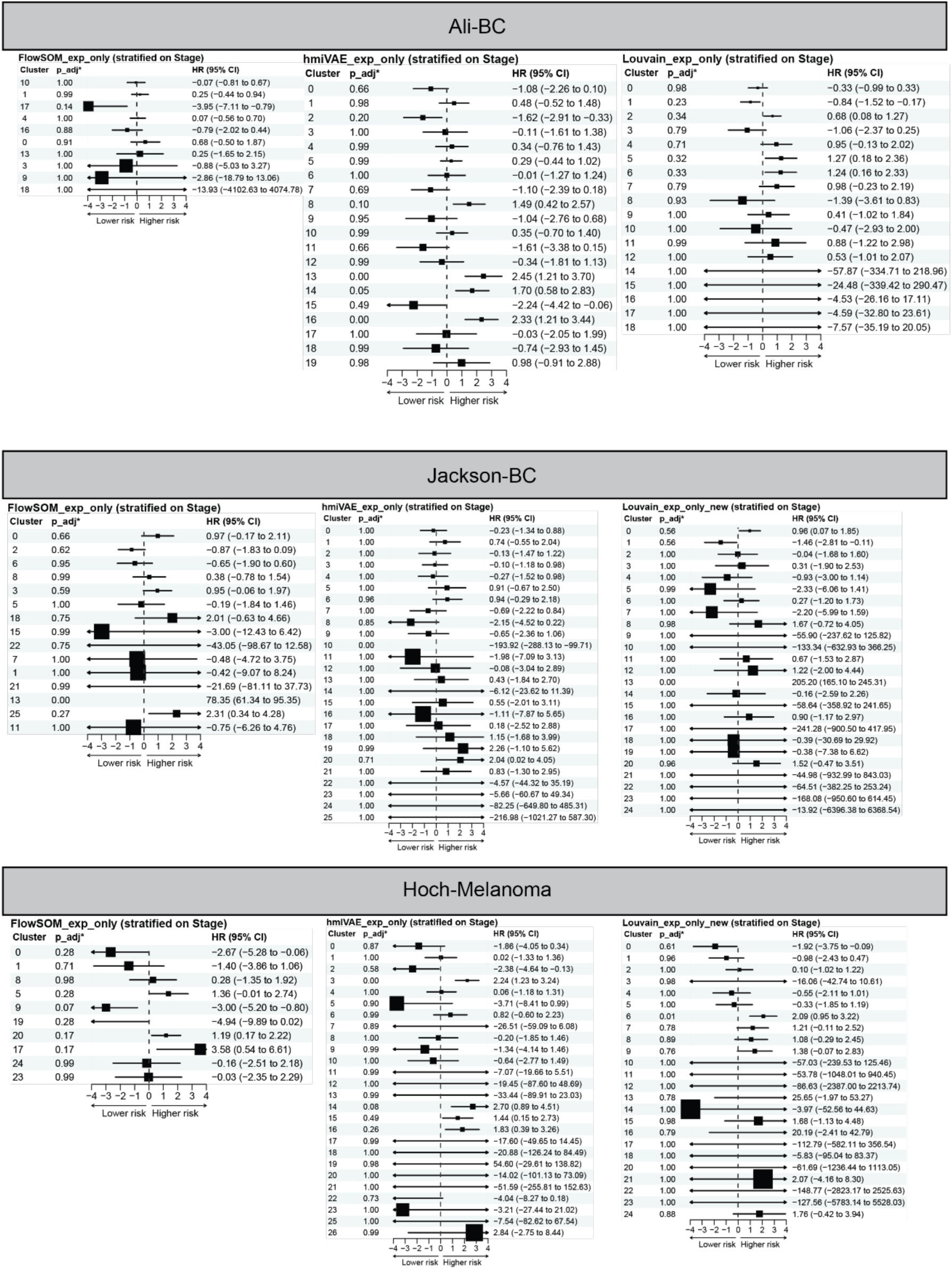
CoxPH model hazard ratios with cluster proportions for clusters from expression feature set for Louvain and FlowSOM and hmiVAE expression embedding clusters for all datasets.

**Supplementary Figure 6.**
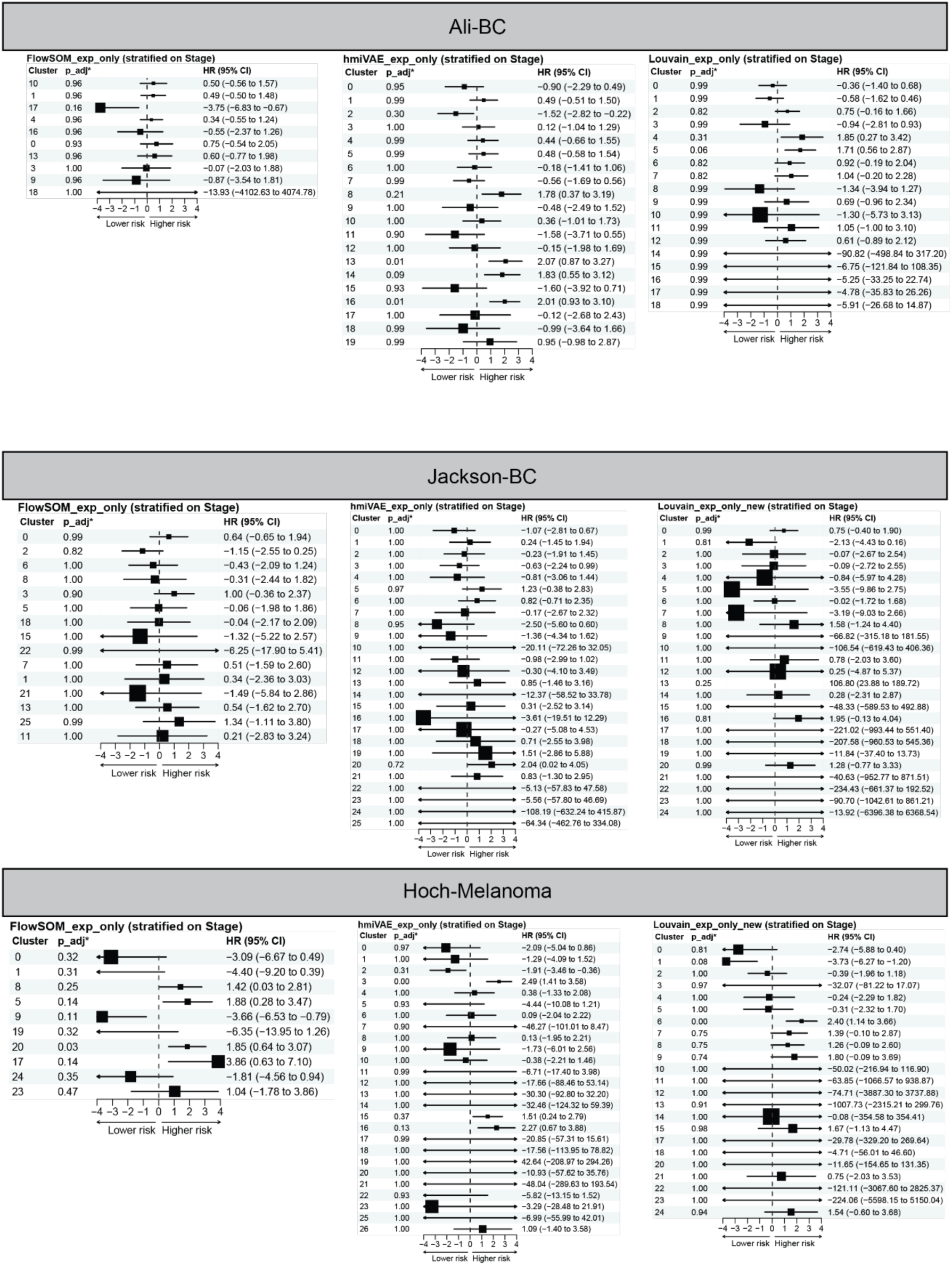
CoxPH model hazard ratios with cluster prevalence per mm^2^ of tissue for clusters from expression feature set for Louvain and FlowSOM and hmiVAE expression embedding clusters for all datasets.

**Supplementary Figure 7.**
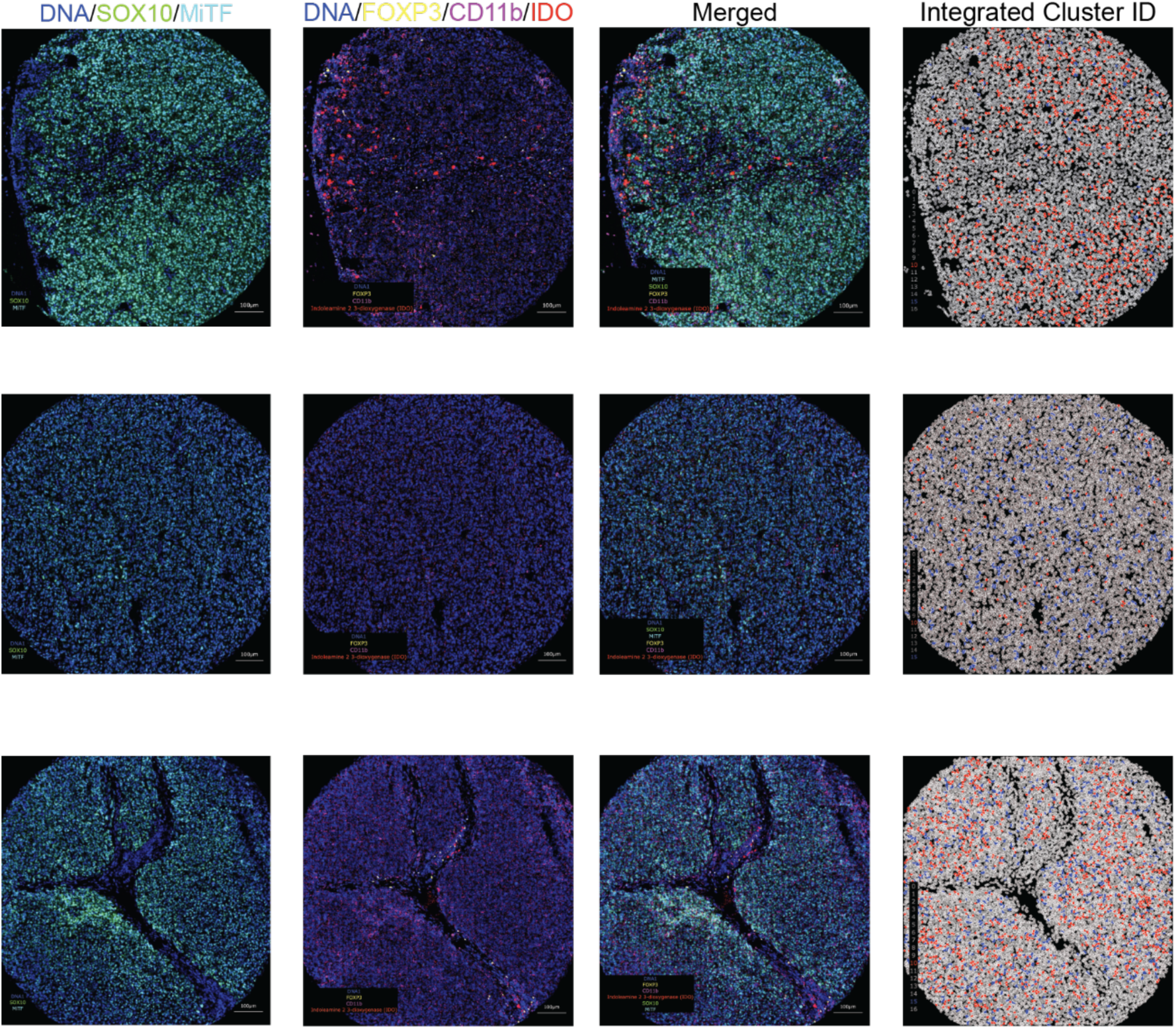
IMC images from Hoch-Melanoma dataset showing patients that had a high number of cells belonging to cluster 10 (top), cluster 15 (middle) or both (bottom). SOX10 and MiTF are tumour cell markers, FOXP3, CD11b and IDO are markers for Tregs, macrophages and T cell suppressive microenvironment, respectively. Integrated cluster IDs, cluster 10 in red and cluster 15 in blue.

**Supplementary Figure 8.**
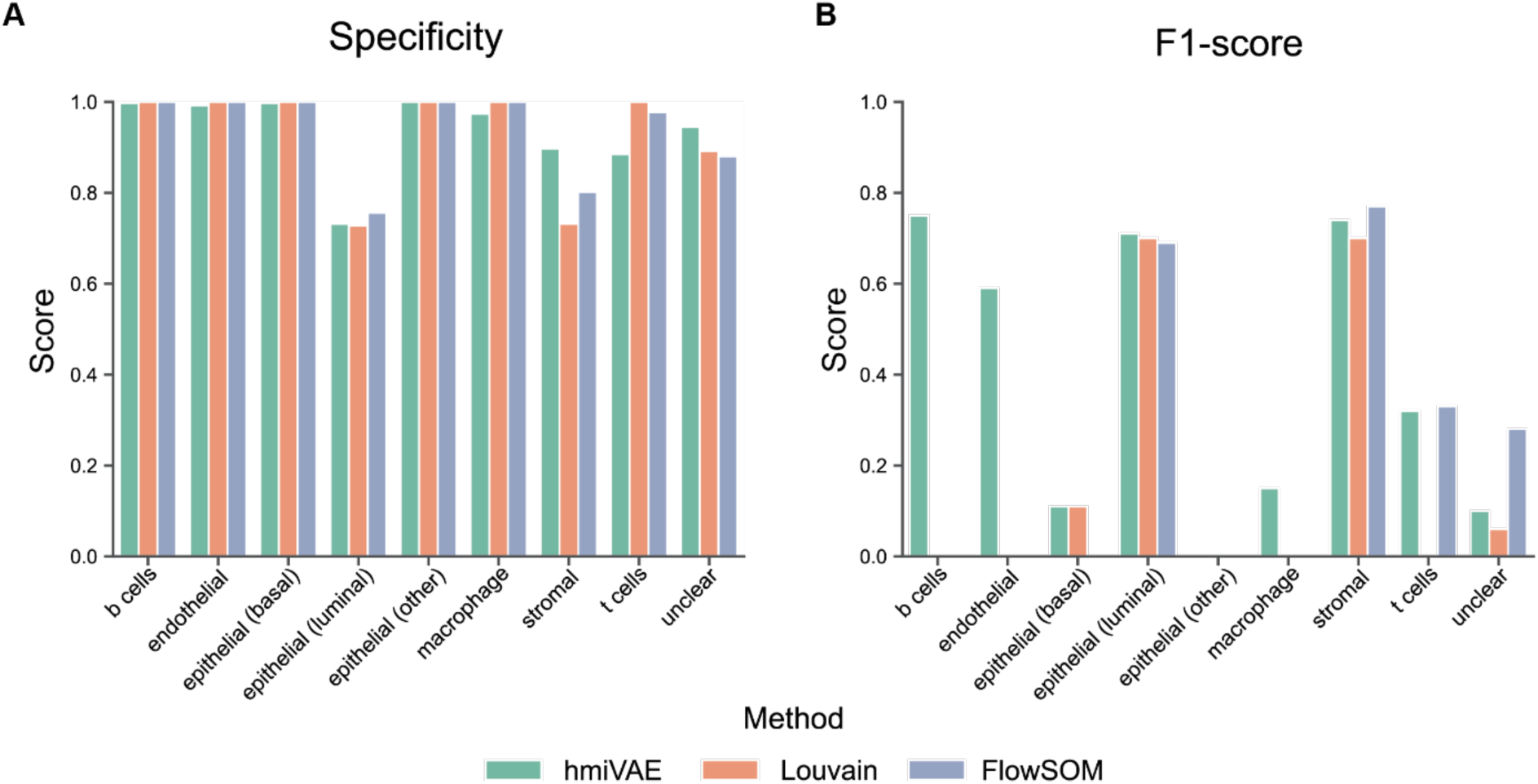
Specificity and F1-Score for all methods in calling the different cell populations when compared to manually annotated ground truth labels of 500 cells.

**Supplementary Figure 9.**
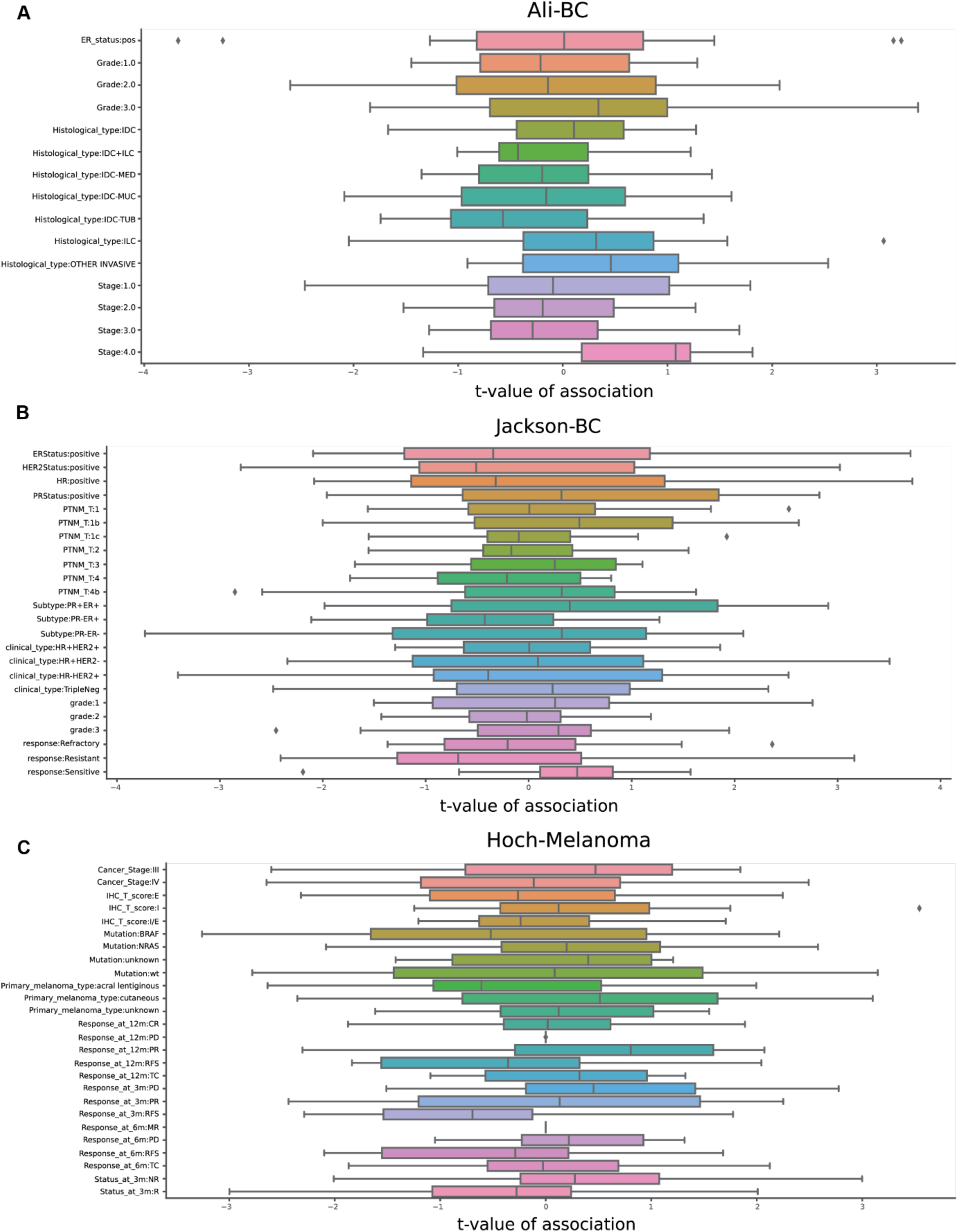
Clinical association of hmiVAE latent space dimensions.

**Supplementary Fig 10.**
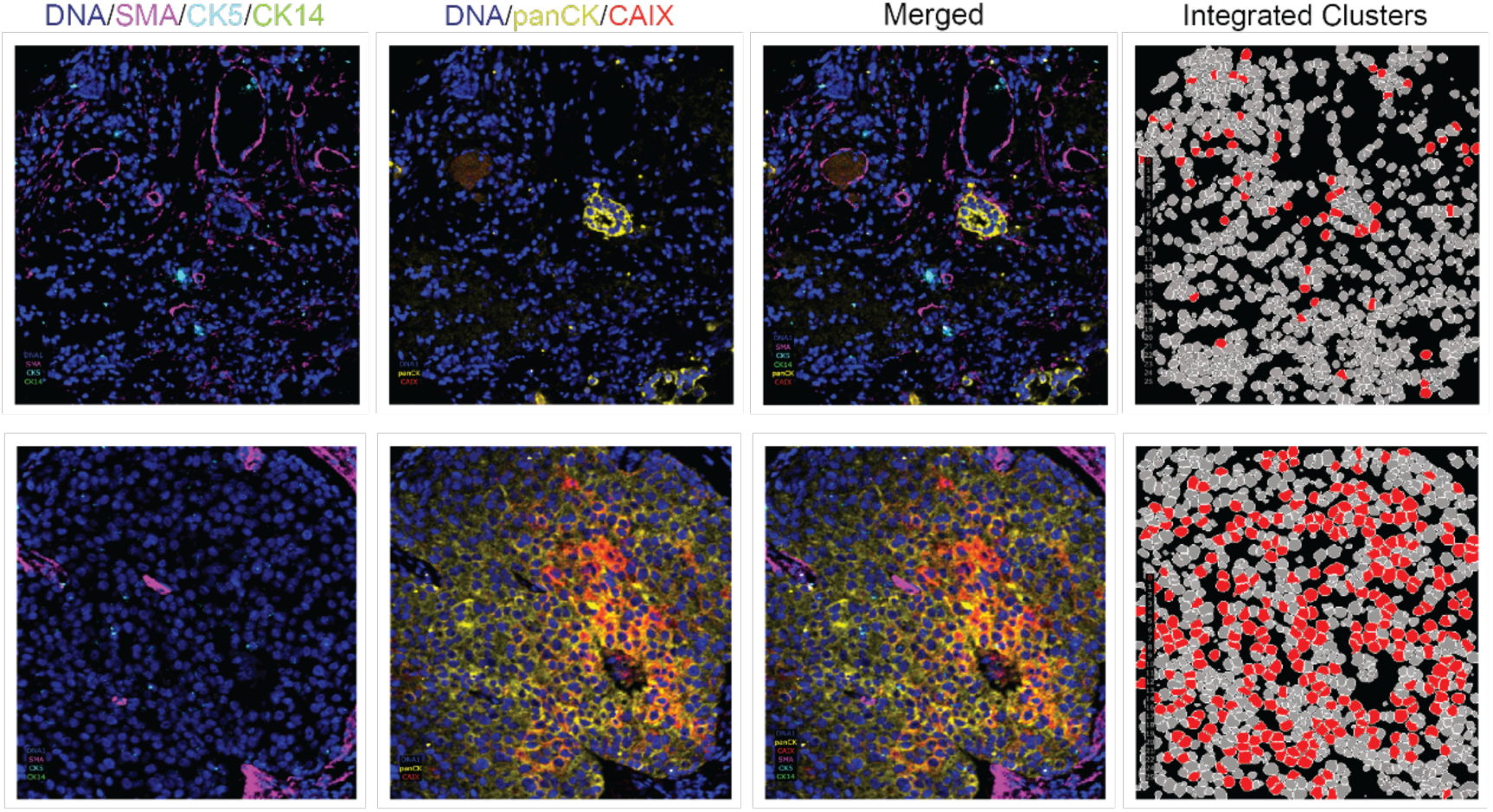
IMC images from the Ali-BC dataset showing marker expression patterns for SMA, CK5, CK14, panCK and CAIX in a patient with low number of cells belonging to cluster 0 (top) and a patient with a high number of cells belonging to cluster 0 (bottom). Cells belonging to integrated cluster 0 are shown in red, while all other clusters are shown in shades of grey.

**Table S1.** Description for all datasets used in this study.

| Cohort | # of Images | # of cells | Panel design | Clinical variables | Tissue |
| --- | --- | --- | --- | --- | --- |
| Jackson-BC | 358 | 802591 | Stromal/Epithelial | HR/HER2 status, Clinical subtype, Cancer stage, Treatment response | Breast |
| Ali-BC | 548 | 536883 | Stromal/Epithelial | ER status, Cancer grade and stage | Breast |
| Hoch-Melanoma | 167 | 989404 | Immune | Cancer stage, Mutation, Primary type, Treatment response | Skin |

**Table S2.** Optimal hmiVAE model hyperparameters for each dataset.

| Cohort | Initialization (random seed) | # of hidden layers | Hidden layer dim. | Latent representation dim. | Beta scheme | Batch size |
| --- | --- | --- | --- | --- | --- | --- |
| Jackson-BC | 123 | 1 | 64 | 20 | Warm-up | 8000 |
| Ali-BC | 1234 | 2 | 32 | 20 | Warm-up | 4000 |
| Hoch-Melanoma | 123 | 1 | 64 | 20 | Warm-up | 16000 |

## References

1. Lin, J.-R. et al. Highly multiplexed immunofluorescence imaging of human tissues and tumors using t-CyCIF and conventional optical microscopes. Elife 7, (2018).

2. Gut, G., Herrmann, M. D. & Pelkmans, L. Multiplexed protein maps link subcellular organization to cellular states. Science 361, (2018).

3. Giesen, C. et al. Highly multiplexed imaging of tumor tissues with subcellular resolution by mass cytometry. Nat. Methods 11, 417–422 (2014).

4. Angelo, M. et al. Multiplexed ion beam imaging of human breast tumors. Nat. Med. 20, 436–442 (2014).

5. Jackson, H. W. et al. The single-cell pathology landscape of breast cancer. Nature 578, 615–620 (2020).

6. Ali, H. R. et al. Imaging mass cytometry and multiplatform genomics define the phenogenomic landscape of breast cancer. Nature Cancer 1, 163–175 (2020).

7. Yousefi, R., Jevdokimenko, K., Kluever, V., Pacheu-Grau, D. & Fornasiero, E. F. Influence of Subcellular Localization and Functional State on Protein Turnover. Cells 10, (2021).

8. Way, G. P. et al. Morphology and gene expression profiling provide complementary information for mapping cell state. Cell Syst 13, 911–923.e9 (2022).

9. Li, D., Ding, J. & Bar-Joseph, Z. Identifying signaling genes in spatial single-cell expression data. Bioinformatics 37, 968–975 (2021).

10. Zidane, M. et al. A review on deep learning applications in highly multiplexed tissue imaging data analysis. Front Bioinform 3, 1159381 (2023).

11. Greenwald, N. F. et al. Whole-cell segmentation of tissue images with human-level performance using large-scale data annotation and deep learning. Nat. Biotechnol. 40, 555–565 (2022).

12. Rahim, M. K. et al. Dynamic CD8+ T cell responses to cancer immunotherapy in human regional lymph nodes are disrupted in metastatic lymph nodes. Cell 186, 1127–1143.e18 (2023).

13. van Maldegem, F. et al. Characterisation of tumour microenvironment remodelling following oncogene inhibition in preclinical studies with imaging mass cytometry. Nat. Commun. 12, 5906 (2021).

14. Geuenich, M. J. et al. Automated assignment of cell identity from single-cell multiplexed imaging and proteomic data. Cell Syst 12, 1173–1186.e5 (2021).

15. Amitay, Y. et al. CellSighter: a neural network to classify cells in highly multiplexed images. Nat. Commun. 14, 4302 (2023).

16. Lee, H.-C., Kosoy, R., Becker, C. E., Dudley, J. T. & Kidd, B. A. Automated cell type discovery and classification through knowledge transfer. Bioinformatics 33, 1689–1695 (2017).

17. Zhang, W. et al. Identification of cell types in multiplexed in situ images by combining protein expression and spatial information using CELESTA. Nat. Methods 19, 759–769 (2022).

18. Tanevski, J., Flores, R. O. R., Gabor, A., Schapiro, D. & Saez-Rodriguez, J. Explainable multiview framework for dissecting spatial relationships from highly multiplexed data. Genome Biol. 23, 97 (2022).

19. Lopez, R. et al. DestVI identifies continuums of cell types in spatial transcriptomics data. Nat. Biotechnol. (2022) doi:10.1038/s41587-022-01272-8.

20. Maseda, F., Cang, Z. & Nie, Q. DEEPsc: A Deep Learning-Based Map Connecting Single-Cell Transcriptomics and Spatial Imaging Data. Front. Genet. 12, 636743 (2021).

21. Lopez, R., Regier, J., Cole, M. B., Jordan, M. I. & Yosef, N. Deep generative modeling for single-cell transcriptomics. Nat. Methods 15, 1053–1058 (2018).

22. Gayoso, A. et al. Joint probabilistic modeling of single-cell multi-omic data with totalVI. Nat. Methods 18, 272–282 (2021).

23. Xu, C. et al. DeepST: identifying spatial domains in spatial transcriptomics by deep learning. Nucleic Acids Res. 50, e131 (2022).

24. Maan, H. et al. Multi-Modal Disentanglement of Spatial Transcriptomics and Histopathology Imaging. Genomics (2025).

25. Flores, M. et al. Deep learning tackles single-cell analysis-a survey of deep learning for scRNA-seq analysis. Brief. Bioinform. 23, (2022).

26. Kingma, D. P. & Welling, M. Auto-Encoding Variational Bayes. arXiv [stat.ML] (2013).

27. Higgins, I. et al. beta-VAE: Learning Basic Visual Concepts with a Constrained Variational Framework. (2016).

28. Simidjievski, N. et al. Variational Autoencoders for Cancer Data Integration: Design Principles and Computational Practice. Front. Genet. 10, 1205 (2019).

29. Lopez, R., Gayoso, A. & Yosef, N. Enhancing scientific discoveries in molecular biology with deep generative models. Mol. Syst. Biol. 16, e9198 (2020).

30. Chow, Y. L., Singh, S., Carpenter, A. E. & Way, G. P. Predicting drug polypharmacology from cell morphology readouts using variational autoencoder latent space arithmetic. bioRxiv 2021.09.02.458673 (2021) doi:10.1101/2021.09.02.458673.

31. Hoch, T., et al. Multiplexed imaging mass cytometry of the chemokine milieus in melanoma characterizes features of the response to immunotherapy. Sci Immunol 7, eabk1692 (2022).

32. Milosevic, V. Different approaches to Imaging Mass Cytometry data analysis. Bioinform. Adv. 3, vbad046 (2023).

33. Yang, L. & Shami, A. On hyperparameter optimization of machine learning algorithms: Theory and practice. Neurocomputing 415, 295–316 (2020).

34. Sønderby, C. K., Raiko, T., Maaløe, L., Sønderby, S. K. & Winther, O. Ladder Variational Autoencoders. arXiv [stat.ML] (2016).

35. Wolf, F. A., Angerer, P. & Theis, F. J. SCANPY: large-scale single-cell gene expression data analysis. Genome Biol. 19, 15 (2018).

36. Vincent D Blondel, Jean-Loup Guillaume, Renaud Lambiotte, Etienne Lefebvre. Fast unfolding of communities in large networks. https://iopscience.iop.org/article/10.1088/1742-5468/2008/10/P10008/pdf (2008) doi:10.1088/1742-5468/2008/10/P10008.

37. Van Gassen, S. et al. FlowSOM: Using self-organizing maps for visualization and interpretation of cytometry data. Cytometry A 87, 636–645 (2015).

38. Warrens, M. J. & van der Hoef, H. Understanding the adjusted Rand index and other partition comparison indices based on counting object pairs. J. Classif. 39, 487–509 (2022).

39. Bao, F. et al. Integrative spatial analysis of cell morphologies and transcriptional states with MUSE. Nat. Biotechnol. (2022) doi:10.1038/s41587-022-01251-z.

40. Madhumita & Paul, S. Capturing the latent space of an Autoencoder for multi-omics integration and cancer subtyping. Comput. Biol. Med. 148, 105832 (2022).

41. Gorringe, K. L. & Fox, S. B. Ductal Carcinoma In Situ Biology, Biomarkers, and Diagnosis. Front. Oncol. 7, 248 (2017).

42. Uyttenhove, C. et al. Evidence for a tumoral immune resistance mechanism based on tryptophan degradation by indoleamine 2,3-dioxygenase. Nat. Med. 9, 1269–1274 (2003).

43. Tie, Y., Tang, F., Wei, Y.-Q. & Wei, X.-W. Immunosuppressive cells in cancer: mechanisms and potential therapeutic targets. J. Hematol. Oncol. 15, 61 (2022).

44. Suzuki, M. & Matsuo, Y. A survey of multimodal deep generative models. Adv. Robot. 36, 261– 278 (2022).

45. T. M. Cover, P. E. H. Nearest Neighbor Pattern Classification. IEEE Trans. Inf. Theory 13, (1966).

46. Maan, H. et al. The differential impacts of dataset imbalance in single-cell data integration. bioRxiv 2022.10.06.511156 (2022) doi:10.1101/2022.10.06.511156.

47. Ramirez Flores, R. O., Lanzer, J. D., Dimitrov, D., Velten, B. & Saez-Rodriguez, J. Multicellular factor analysis of single-cell data for a tissue-centric understanding of disease. Elife 12, (2023).

48. Liu, C. C. et al. Robust phenotyping of highly multiplexed tissue imaging data using pixel-level clustering. Nat. Commun. 14, 4618 (2023).

49. He, F., Liu, T. & Tao, D. Control batch size and learning rate to generalize well: Theoretical and empirical evidence. Adv. Neural Inf. Process. Syst. 1141–1150 (2019).

50. Steck, H., Ekanadham, C. & Kallus, N. Is Cosine-Similarity of Embeddings Really About Similarity? arXiv [cs.IR] (2024).

51. Carstens, J. L. et al. Spatial multiplexing and omics. Nat. Rev. Methods Primers 4, 1–19 (2024).

52. Curtis, C. et al. The genomic and transcriptomic architecture of 2,000 breast tumours reveals novel subgroups. Nature 486, 346–352 (2012).

53. van der Walt, S. et al. scikit-image: image processing in Python. PeerJ 2, e453 (2014).

54. Traag, V. A., Waltman, L. & van Eck, N. J. From Louvain to Leiden: guaranteeing well-connected communities. Sci. Rep. 9, 5233 (2019).

55. Pedregosa, F. et al. Scikit-learn: Machine Learning in Python. J. Mach. Learn. Res. abs/1201.0490, (2011).

56. Davidson-Pilon, C. lifelines: survival analysis in Python. J. Open Source Softw. 4, (2019).

